# Inhibition governs preference encoding in medial prefrontal cortex pyramidal neurons during a binary social choice in mice

**DOI:** 10.1101/2025.11.16.688668

**Authors:** Renad Jabarin, Paritosh Jaiswal, Shai Netser, Shlomo Wagner

## Abstract

Social decision-making requires the brain to evaluate competing options and select appropriate actions. The medial prefrontal cortex (mPFC) is implicated in social cognition and value-based decision-making, yet how it encodes the relative value of competing social choices remains unclear. Here, we used fiber photometry, optogenetics, and projection-specific recordings in mice performing four binary social discrimination tasks to examine how mPFC pyramidal neurons encode social choice. We found that these neurons showed marked inhibition during bouts toward the preferred stimulus. This inhibition was specific to transitional bouts, when animals switched between stimuli, and was absent during repeated, non-transitional bouts. Negative calcium transients predicted subsequent investigation of the preferred stimulus, indicating a functional role in guiding choice. Importantly, this inhibition encoded relative stimulus value rather than identity. Optogenetic activation of mPFC pyramidal neurons during investigation induced immediate avoidance, yet paradoxically promoted persistent re-engagement with the same option through repetitive non-transitional bouts. Projection-specific recordings further revealed differential recruitment of mPFC neurons targeting the nucleus accumbens and basolateral amygdala across tasks. Together, these findings identify inhibition of mPFC pyramidal neurons as a neural signature of stimulus preference, revealing a principle by which the mPFC dynamically guides social choice.

## Introduction

Social interactions are fundamental to the survival and well-being of many species, including humans. These interactions comprise a continuous stream of decisions that require an individual to evaluate available social partners, interpret their intentions, and select appropriate actions [1, 2]. Thus, social decision-making is a complex cognitive process that relies on the brain’s ability to assign value to different social stimuli and to flexibly adapt behavior in response to a constantly changing social landscape [3]. A breakdown in this critical capacity can lead to profound social deficits, as seen in a variety of neuropsychiatric and neurodevelopmental disorders, most notably autism spectrum disorder (ASD) [4, 5].

The medial prefrontal cortex (mPFC) has emerged as a critical hub in the neural circuitry governing social cognition and value-based decision-making [6, 7]. Extensive research in both humans and animal models has identified the mPFC as a critical regulator of a wide range of social functions, including recognition of conspecifics, interpretation of social cues, regulation of approach/avoidance behavior [8–10] and integration of emotional and contextual cues [11]. This brain region is not a homogeneous structure; it comprises at least two functionally distinct subregions along its dorsoventral axis: the prelimbic cortex (PrL) and the infralimbic cortex (IL) [12, 13]. The PrL has been consistently implicated in the active expression of goal-directed and approach behavior, including social recognition and investigation, and plays a key role in guiding adaptive social choices [8, 14]. The IL has been more closely associated with the suppression of ongoing behavior, fear extinction, and inhibitory control over reward-seeking, and has been proposed to mediate avoidance and the flexible updating of behavioral responses [14–16]. Despite their often opposing functional profiles, both subregions are anatomically interconnected with limbic and striatal circuits, including the nucleus accumbens (NAc) and the basolateral amygdala (BLA), through which they exert top-down control over motivated behavior [13, 17].

While it is well established that the mPFC encodes context-dependent value in non-social operant tasks involving explicit rewards or costs [18], it remains unclear whether similar principles govern the more fluid and spontaneous nature of social interactions. In natural settings, the ‘value’ of a social partner is not fixed but rather dynamically shaped by the presence of other individuals, the subject’s internal state, and the broader social environment [19, 20]. This concept of relative valuation is a hallmark of flexible and adaptive decision-making. However, a significant gap remains in our understanding of how the mPFC translates the relative value of social options into a concrete behavioral choice [21, 22]. Specifically, it is still unknown how the mPFC dynamically encodes the relative value of competing social and non-social options and how this evolving representation translates into behavioral choice [12, 23].

Here, we combine fiber photometry from mPFC pyramidal neurons with a battery of four distinct social discrimination tasks in mice, each requiring a choice between two stimuli of varying motivational value, to show that the mPFC encodes the relative value of social choices through selective inhibition: calcium signals in pyramidal neurons are profoundly inhibited during transitional bouts toward the preferred stimulus, distinguishing it from increased activity characterizing investigation of both the non-preferred stimulus and empty chambers. The fact that this differential activity was observed specifically during transitional bouts between stimuli suggests that it has a role in the decision to engage with a new option. Accordingly, we found that negative calcium transients in the mPFC were predictive of subsequent investigation of the preferred stimulus. Furthermore, social fear conditioning, which selectively inverted the behavioral preference for a previously neutral social stimulus, also inverted its neural signature, converting the inhibitory response into excitation, consistent with a caution signal that reflects aversive rather than positive valuation. Distinct mPFC projections to the NAc and BLA were differentially engaged during various social decisions, indicating that valuation and emotional discrimination are mediated through segregated downstream circuits. Finally, optogenetic activation of mPFC pyramidal neurons during stimulus investigation, despite eliciting an immediate avoidance, was sufficient to bias choice by increasing the preference for the stimulated option via repetitive bouts of investigation. Together, these findings identify a neural principle underlying social choice encoding in the mPFC and demonstrate how mPFC circuits transform social information into adaptive decisions.

## Results

### I. Behavioral characterization of social choice across multiple binary discrimination tasks

To establish a behavioral framework for interpreting neural activity in the mPFC, we first assessed how subject mice behave when faced with a binary choice between distinct stimuli with varying motivational value. To that end, adult males were tested in four different discrimination tasks requiring them to choose between two distinct stimuli.

In the Social Preference (SP) task, subject mice were exposed to a novel age- and sex-matched conspecific (social stimulus) on one side of the arena and an inanimate object on the other side (Fig. 1A), and exhibited a higher preference to investigate the social stimulus over the object, as reflected by the investigation time (Fig. 1B-C) and mean investigation bout duration (Fig. 1D) for each of the stimuli. In the Sex Preference (SxP) task, subjects preferred to investigate a female over a male conspecific (Fig. 1F-I). In the Emotional State Preference for stress (ESPs) task, subjects preferred a stressed over a naïve conspecific (Fig. 1K-N). Lastly, in the Social vs. Food Preference (SvFP) task, subjects displayed a significant preference for a conspecific over food (Fig. 1P-S). While the four tasks represent distinct social contexts, they all exhibit a similar level of stimulus preference, as reflected by the relative discrimination index (RDI; Fig. S1A) of the preferred stimulus in each task. Notably, all four tasks shared one type of stimulus – a novel naïve male, which was the preferred stimulus in the SP and SvFP tasks, but was the non-preferred stimulus in the SxP and ESPs tasks.

**Figure 1.**
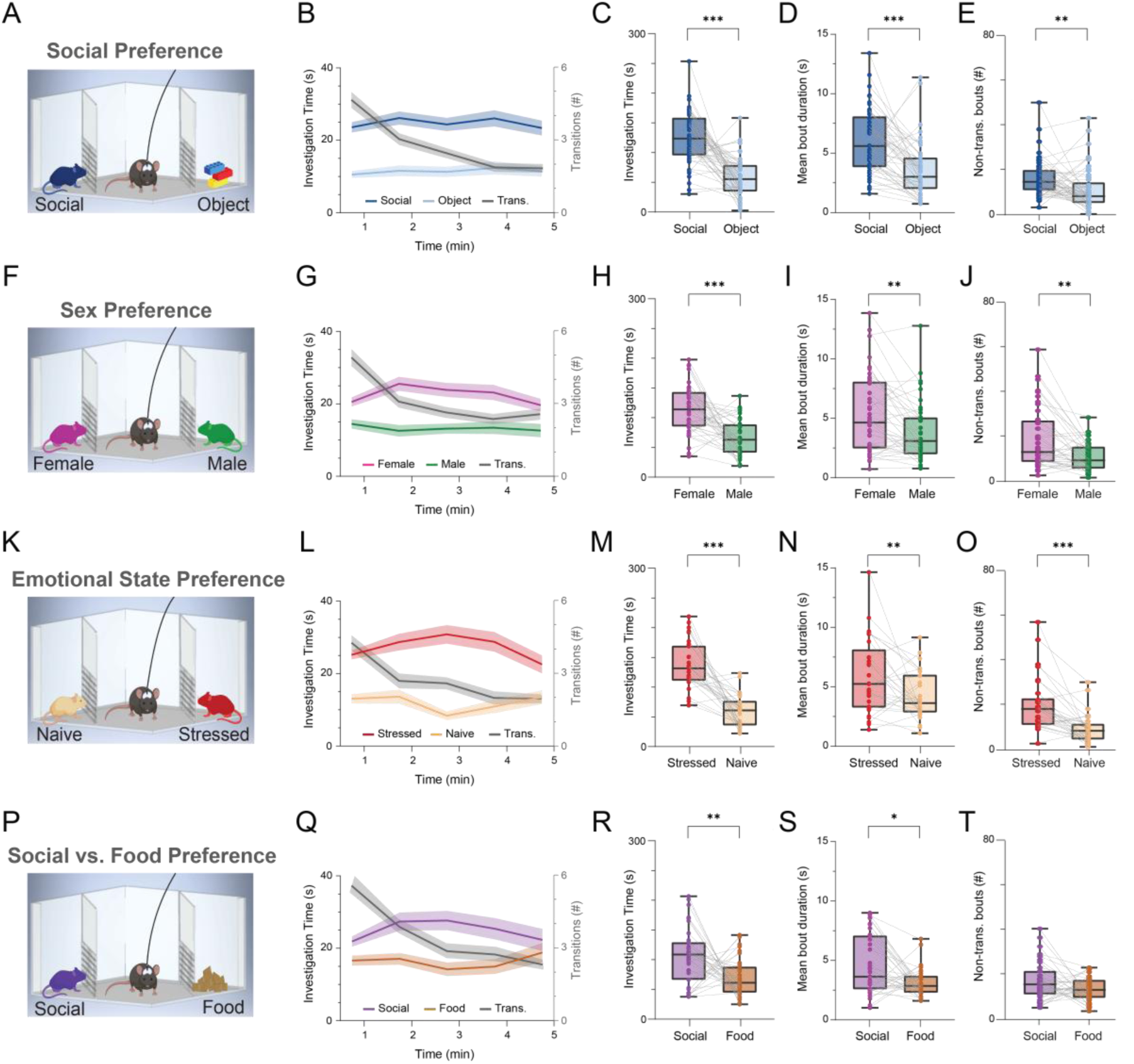
Behavioral characterization of social choice across multiple binary discrimination tasks. **A.** Schematic representation of the SP task. **B.** Mean time (±SEM) dedicated by subject mice (n=49 sessions from 21 animals) for investigating the animal stimulus (dark blue) or the object (light blue), and mean sum of transitional bouts between the two stimuli (±SEM, gray line) across the 5 min encounter period of the SP task in (1-min bins). **C.** Median (in box plot*) time of investigating the social stimulus and the object. Each grey line connects data points of the same session. **D.** Median (in box plot^&^) duration of all investigation bouts with the social stimulus and the object. **E.** Median (in box plot^&^) sum of non-transitional bouts with the social stimulus and the object. **F-J.** As in **A-E**, for the SxP task (n=40 sessions from 14 animals). **K-O.** As in **A-E**, for the ESPs task (n=30 sessions from 14 animals). **P-T.** As in **A-E**, for the SvFP task (n=35 sessions from 14 animals). \*\*\**p*<0.001, paired samples t-test in **C**, **H**, **M** and **R**. *\*p<0.05*, \*\**p*<0.01, \*\*\**p*<0.001, Wilcoxon signed-rank test in **D-E**, **I-J**, **N-O**, and **S-T**. ^&^ Box plot represents 25 to 75 percentiles of the distribution, while the bold line is the median of the distribution. Whiskers represent the smallest and largest values in the distribution.

The four tasks also shared a similar level of social propensity, as reflected in the total investigation time (Fig. S1B) and a similar distance traveled during each task (Fig. S1C). Thus, they appear to induce a comparable level of social motivation and arousal in the subject animals, and differ primarily in the type of choice, dictated by the identity of the presented stimuli.

We then differentiate between two types of investigation bouts presumably differing in the type of decision preceding them: transitional vs. non-transitional. Transitional bouts occur when the subject animal investigates one stimulus following a previous bout with the other stimulus. As we previously described [24], transitional bouts were more abundant at the early stage of each of the four tasks, with no apparent correlation with the investigation time (Fig. 1B,G,L,Q, grey line). They reflect the animal’s decision to stop investigating one stimulus and start investigating the other stimulus, thus performing a behavioral shift. In contrast, non-transitional bouts occur when the subject animal investigates one stimulus following a previous bout with the same stimulus, hence reflecting the sustained tendency of the animal to pursue the same stimulus.

We found that in all four tasks, mice exhibited a greater number of non-transitional bouts compared to transitional bouts (Fig. S1D). Moreover, we found a significantly higher number of non-transitional bouts for the preferred stimulus over the non-preferred stimulus in all tasks, except for the SvFP, where only a trend was observed (Fig. 1E,J,O,T). Thus, mice exhibit a persistent tendency to investigate the preferred stimulus once it has been selected. Altogether, these findings suggest that and that social decision-making involves both sustained engagement and selective switching between options and that transitional and non-transitional investigation bouts may be used as measurements of these two types of social decision-making.

### II. Activity of mPFC pyramidal neurons discriminates between transitional and non-transitional bouts

To examine the activity of mPFC CaMKII*α*-expressing pyramidal neurons during the four social discrimination tasks, we employed fiber photometry, as previously described [25]. For that, we injected the mPFC with AAV viral vectors to express the genetically encoded calcium indicator GCaMP6f under the control of the CaMKII*α* promoter (Fig. 2A). We then recorded calcium signals from subject mice during the experiments shown in Fig. 1, with each session comprising a 10-minute recording of calcium signals: five minutes before the introduction of stimuli into the empty chambers (pre-encounter stage) and five minutes after it (encounter stage, see timeline in Fig. 2B). Example traces of normalized calcium signal aligned with the onset of non-transitional and transitional bouts performed during that session, as well as mean traces for the encounter phase for each of the four tasks are shown in Fig. S2A-H.

**Figure 2.**
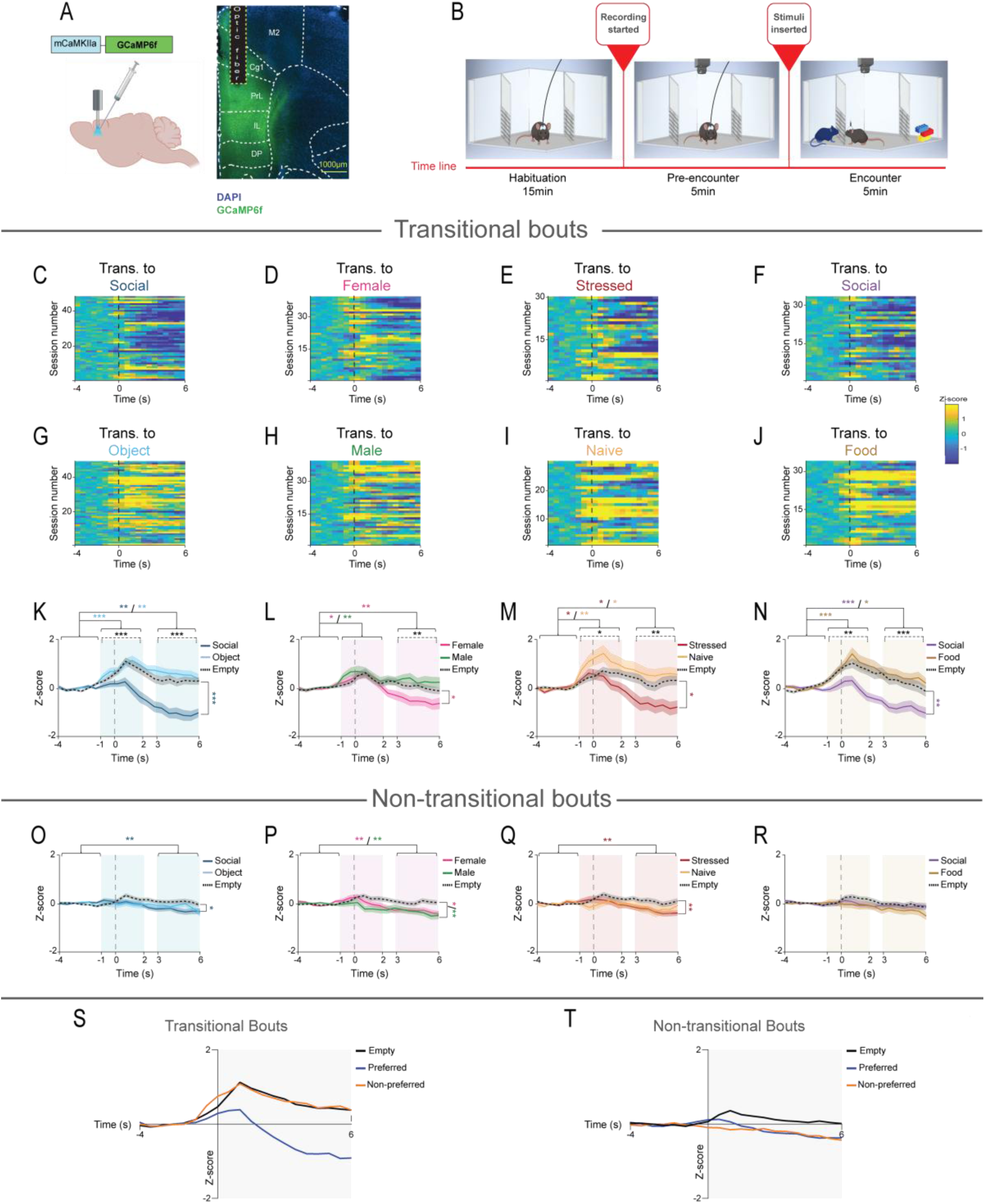
Activity of mPFC pyramidal neurons discriminates between transitional and non-transitional bouts. **A.** On the left, a schematic representation of the GCaMP6-expressing AAV vector injection followed by optic fiber implant in the mPFC. On the right, a fluorescent micrograph of an AAV vector-infected mPFC slice, showing GCaMP6-expression (green) and optic fiber location (dashed line). **B.** Timeline for a typical session of behavioral testing with fiber photometry recording. **C.** Heat-map (see color code to the right of **F**) of the z-scored calcium signals at the beginning (first six seconds) of bout, averaged across all transitional bouts towards the stimulus animal, shown for all sessions of the SP task (n=49 sessions, with each row corresponding to a single session), using 0.5 sec bins. Time ‘0’ represents the beginning of the bout. **D-J.** As in **C,** for the preferred (female, stressed, and social) stimuli and non-preferred stimuli (object, male, naïve, and food), respectively, in each task (SxP; n_SxP_=40; n_ESPs_=30; n_SvFP_=35). **K.** Superimposed traces of the mean (±SEM) z-scored calcium signals shown in **C and G**, averaged across all SP sessions. The dark blue and light blue trace lines represent the z-scored calcium signals in transitional bouts to the social stimulus and the object, respectively, during the encounter stage of the SP sessions. Dashed black traces are for transitional bouts to empty chambers during the pre-encounter stage. Blue-shaded areas represent two distinct time windows of the event, which are used for statistical analysis. Asterisks, representing statistical comparisons between the stimuli and across time windows, are color-coded according to the type of comparison at the top of the graph. Blue and light blue asterisks represent a significant difference in social and object z-scored signals, respectively, during the specific time window relative to baseline. Black asterisks on dashed lines represent a difference in responses between the two stimuli during the same time window. Asterisks on the right side of the graph represent statistical differences for social and object z-scored signals in the encounter stage of the session, during the 3-6s time window, relative to the pre-encounter stage, and are color-coded according to the stimulus with the significant difference. A significant effect was found for stimulus (*p<0.001*), time (*p<0.001*), and stimulus x time interaction (*p<0.001*) by two-way MM ANOVA. **L-N.** As in **K**, for transitional bouts in the SxP, ESPS, and SvFP tasks, respectively. In all three tasks, a significant effect was found for stimulus (*p<0.05, p<0.01, p<0.001*, respectively), time (*p<0.001*), and stimulus x time interaction (*p<0.01* in SxP and ESPs*; p<0.001* in SvFP) by two-way MM ANOVA. **O-R.** As in **K**, for non-transitional bouts in the SP, SxP, ESPs, and SvFP tasks, respectively. A significant effect was found for time (*p<0.01* in SP, *p<0.001* in SxP, *p<0.01* in ESPs, and *p<0.05* in SvFP) by two-way MM ANOVA. **S.** Mean trace for the z-scored activity during all transitional bouts from the pooled sessions of the four tasks during the pre-encounter stage (in black), transitional bouts to the preferred stimulus (in blue), and transitional bouts to the non-preferred stimulus (in orange). Please note the difference between the inhibitory response in the trace for transitional bouts to the preferred stimulus, compared to the traces for transitional bouts in the pre-encounter stage and to the less-preferred stimulus. **T.** As in **S**, for non-transitional bouts. \**p*<0.05, \*\**p*<0.01, \*\*\**p*<0.001, *post-hoc* paired (within stimulus differences in time) or independent (differences between stimuli within time windows, or differences between pre-encounter and encounter) t-test with Holm-Šídák correction for multiple comparisons following detection of main effects by ANOVA.

We used z-score analysis to normalize and average all the instantaneous signals recorded during the two types of behavioral events -namely, transitional and non-transitional bouts-separately. We then analyzed the averaged z-scored calcium response for each behavioral event across all sessions of the same task, as previously described by us [25]. We first determined the responses of mPFC pyramidal neurons during transitional and non-transitional bouts of investigation conducted toward the two empty chambers during the pre-encounter phase of each session. No behavioral preference was found for one empty chamber over the other in any of the four tasks (Fig. S3A-D). Still, we found that both non-transitional (in SP, SxP, and ESPs tasks; Fig. S3E-F) and transitional bouts (in all four tasks; Fig. S3G-H) elicited an immediate increase in the recorded signal (mainly during -1 to 2s relative to the beginning of the bout), with this excitatory response being more robust in transitional compared to non-transitional bouts (-1 to 2s: t(45)=-5.35, p<0.001; 3 to 6s: t(45)=-1.572, p=0.117, Independent t-test).

We then analyzed the responses of mPFC pyramidal neurons during the encounter stage of the various sessions, when subject mice investigated stimulus-containing chambers. Importantly, we compared these responses with the signals recorded during transitional and non-transitional bouts toward the empty chambers during the pre-encounter stage. We found that transitional bouts in the encounter phase elicited differential neural responses between the two stimuli, with the preferred stimulus eliciting a significant inhibition while the non-preferred one eliciting excitation. Nevertheless, there was no difference between the excitatory response toward the non-preferred stimulus and the response toward the empty chambers during the pre-encounter stage, while the inhibitory response toward the preferred stimulus showed a significant difference from the empty chamber response in all tasks (Fig. 2C-N). In contrast, we found that across all four tasks, non-transitional bouts during the encounter stage did not show any difference in the response between the stimuli (Fig. O-R; Fig. S4A-H). These results, which are further apparent when all four tasks are averaged (Fig. 2S-T), suggest that a differential response between the two stimuli occur in mPFC pyramidal neurons only during transitional investigation bouts, but not during non-transitional bouts. They also suggest that it is the preferred stimulus that elicit an inhibitory response during transitional bouts, while the excitation observed during non-preferred stimulus investigation is a generic response during chamber investigation.

### III. Context-dependent encoding of relative stimulus value in the mPFC

Our results so far show that mPFC CaMKII*α*^+^ neurons are differentially responsive to the distinct stimuli during investigation bouts following a behavioral shift, where the animal leaves one stimulus in order to investigate the other one, while being less (and uniformly) responsive during repeated investigation bouts. Yet, the differential response during transitional bouts may be sensitive to the absolute appeal of the stimulus, determined by its identity, or to its relative appeal, determined by the other option (stimulus) presented in the same context.

As motioned above, all four tasks used by us shared a common stimulus - a naïve male conspecific. However, the preference for this stimulus differed between the tasks: it was the preferred stimulus in the SP and SvFP tasks, while in SxP and ESPs tasks it was the non-preferred stimulus, as shown by the common stimulus discrimination index values of the various tasks (Fig. 3A). Despite the identical nature of this common stimulus across the four tasks, mPFC pyramidal neurons activity during transitional bouts toward this stimulus depended on whether the stimulus was the preferred or the non-preferred in that given task (Fig. 3B-C). Specifically, neural activity during transitional bouts toward the common stimulus was mostly inhibitory in sessions where the common stimulus was the preferred stimulus (Common stimulus discrimination index > 0), while in sessions where it was the non-preferred stimulus (Common stimulus discrimination index < 0) the neural activity was excitatory (Fig. 3D). This difference was specific for transitional bouts and was not detected during non-transitional bouts (Fig. 3E-F).

**Figure 3.**
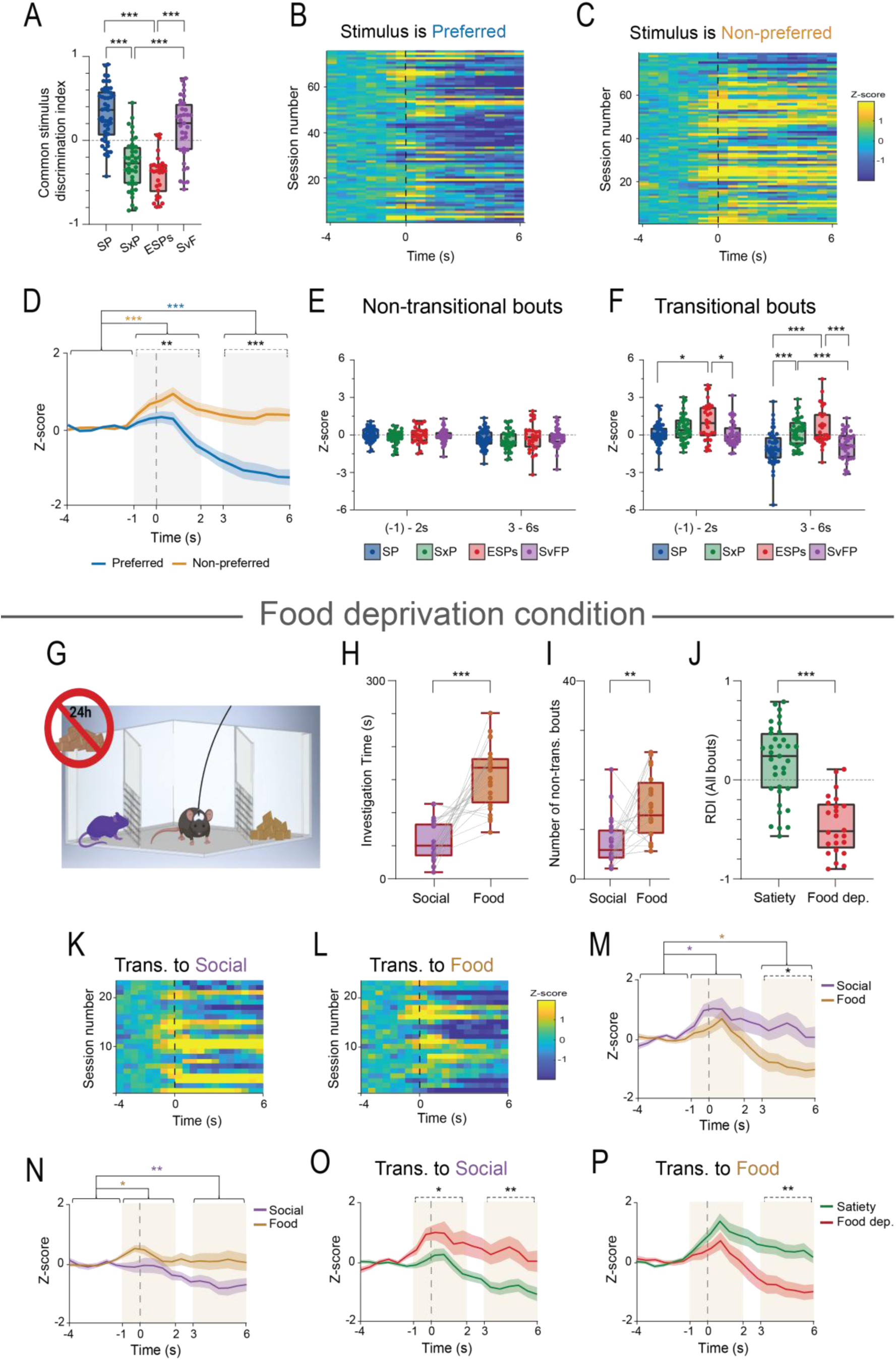
Context-dependent encoding of relative stimulus value in the mPFC. **A.** Median (in box plot^&^) relative discrimination index (RDI) for the common stimulus in each of the four tasks. A significant effect was found for task (*p<0.001*) by one-way ANOVA. **B.** Heat-map (see color code to the right of **C**) of the average z-scored calcium signals at the beginning (first six seconds) of transitional bouts to the preferred stimulus, using 0.5-s bins, pooled from all four tasks. Time ‘0’ represents the beginning of the bout. **C.** As in **B**, for transitional bouts to the non-preferred stimulus, pooled from all four tasks. **D.** Superimposed traces of the mean (±SEM) z-scored calcium signals shown in **B-C**, averaged across all sessions. Gray-shaded areas represent two distinct time windows of the event, which were used for the statistical analysis. A significant effect was found for stimulus (*p<0.001*), time (*p<0.001*), and stimulus x time interaction (*p<0.001*) by two-way MM ANOVA. **E.** Median (in box plot^&^) z-score of the calcium signal during non-transitional bouts to the common stimulus in each of the four tasks, divided into two time-windows: -1 to 2s and 3 to 6s relative to the bout onset. A significant effect was found for time (*p<0.*001) by two-way MM ANOVA. **F.** As in **E**, for transitional bouts. A significant effect was found for stimulus (*p<0.001*), time (*p<0.001*), and stimulus x time interaction (*p<0.001*) by two-way MM ANOVA. **G.** Schematic representation of the SvFP task following 24 hours of food deprivation. **H.** Median (in box plot^&^) time dedicated by subject animals to investigate the social stimulus (in purple) and the food (in brown, n=24 sessions from 14 animals) during the 5 min encounter stage of the SvFP-food deprivation task. Each grey line connects data points of the same session. **I.** Median (in box plot^&^) sum of non-transitional bouts to the social stimulus and the food during the 5 min encounter stage of the task. **J.** Median (in box plot^&^) relative discrimination index (RDI) between the social and food stimuli during the 5 min encounter stage of the two conditions of the SvFP tasks (satiety and food deprivation). **K.** Heat-map (see color code to the right of **L**) of the z-scored calcium signals at the beginning (first six seconds) of transitional bouts, averaged across all transitional bouts, to the social stimulus during the 5 min encounter stage of all sessions of the SvFP-food deprivation task. **L.** As in **K**, for transitional bouts to the food stimulus. **M.** Superimposed traces of the mean (±SEM) z-scored calcium signals shown in **K-L,** averaged across all SvFP-food deprivation sessions. Beige-shaded areas represent two distinct time windows of the event, which are used for the statistical analysis. A significant effect was found for stimulus (*p<0.05*), time (*p<0.001*), and stimulus x time interaction (*p<0.05*) by two-way MM ANOVA. **N.** As in **M**, for non-transitional bouts. A significant effect was found for stimulus (*p<0.01*) by two-way MM ANOVA. **O.** Superimposed traces of the mean (±SEM) z-scored calcium signals for transitional bouts to the social stimulus in the two conditions of the SvFP task. **P.** As in **O**, for transitional bouts to the food stimulus. *\*p<0.05*, \*\**p*<0.01, \*\*\**p*<0.001, *post-hoc* paired or independent t-test with Holm-Šídák correction for multiple comparisons following detection of main effects by ANOVA in **A, D-F**, **M-N**. *\*p<0.05*, \*\**p*<0.01, independent t-test with Holm-Šídák correction for multiple comparisons in **O-P**. ^&^ Box plot represents 25 to 75 percentiles of the distribution, while the bold line is the median of the distribution. Whiskers represent the smallest and largest values in the distribution.

To directly test whether the observed inhibition of mPFC pyramidal neurons in transition bouts is indeed related to the relative appeal of the stimulus rather than its identity, we examined how the neural activity of mPFC pyramidal neurons changes when stimuli identity is held constant while their relative appeal is altered. To that end, we recorded animals performing the SvFP task following 24 hours of food deprivation (Fig. 3G). As expected, in this condition the animals exhibited a strong preference for the food over the social stimulus, as reflected by the time dedicated to investigate each stimulus and the number of non-transitional bouts (Fig. 3H-I). Accordingly, social stimulus RDI values were significantly different between satiety and food deprivation, with a positive median during satiety and a negative median during food deprivation (Fig. 3J). This change in preference was accompanied by a parallel change in neural activity, with transitional bouts to food exhibiting significantly inhibitory responses compared to the social stimulus (Fig. 3K-M), while no differences were found during non-transitional bouts (Fig. 3N). More specifically, the “flipped” responses during transitional bouts to the same stimulus, whether food or social, were opposite between satiety, when the social stimulus was preferred, and food deprivation, when food was the preferred stimulus (Fig. 3O-P). These results further demonstrate that mPFC neurons’ differential response during transitional investigation bouts depends on the relative appeal of the stimulus, with an inhibitory response elicited specifically by the preferred stimulus.

Notably, a histological mapping of the implanted optic fiber tip locations (Fig. S5A) revealed that recordings were obtained from both PrL (44%) and IL (66%) (Fig. S5B). To examine whether the recorded calcium signals correlated with the precise dorsoventral location of the optic fiber, we calculated the Pearson correlation coefficient between the averaged z-scored fluorescence signal and the fiber depth for each task, stimulus, and time window, analyzing transitional and non-transitional bouts separately. This analysis revealed no significant correlations between optic fiber depth and the recorded neural activity during either non-transitional (Fig. S5C-F) or transitional bouts (Fig. S5G-J) in any of the four tasks. Furthermore, even when the data were pooled across all tasks to compare responses to preferred and non-preferred stimuli, no significant correlations were found for non-transitional or transitional bouts (Fig. S5K-L). These results suggest that, within the context of our behavioral paradigms, the observed mPFC pyramidal cell activity -notably the inhibition during transitional bouts to the preferred stimulus- is a general phenomenon across both the PrL and IL regions.

The relationship between inhibited activity of mPFC pyramidal neurons and social preference was further assessed by calculating the probability of social investigation following negative or positive changes in calcium activity (Fig. S6A-K). This analysis revealed that negative calcium transients are predictive of the subjects’ behavior in all four tasks, given that they were followed by a statistically significant increased probability of investigation behavior specifically toward the preferred stimulus. (Fig. 4A-P,Q). Moreover, this increase in probability during the encounter was significantly higher than that recorded during the pre-encounter stage, suggesting that the negative transients are associated with the investigation of the preferred stimulus. In contrast, no such changes of significantly higher probability to investigate a certain stimulus compared to pre-encounter were found for positive transients (Fig. 4R; Fig. S7A-H).

**Figure 4.**
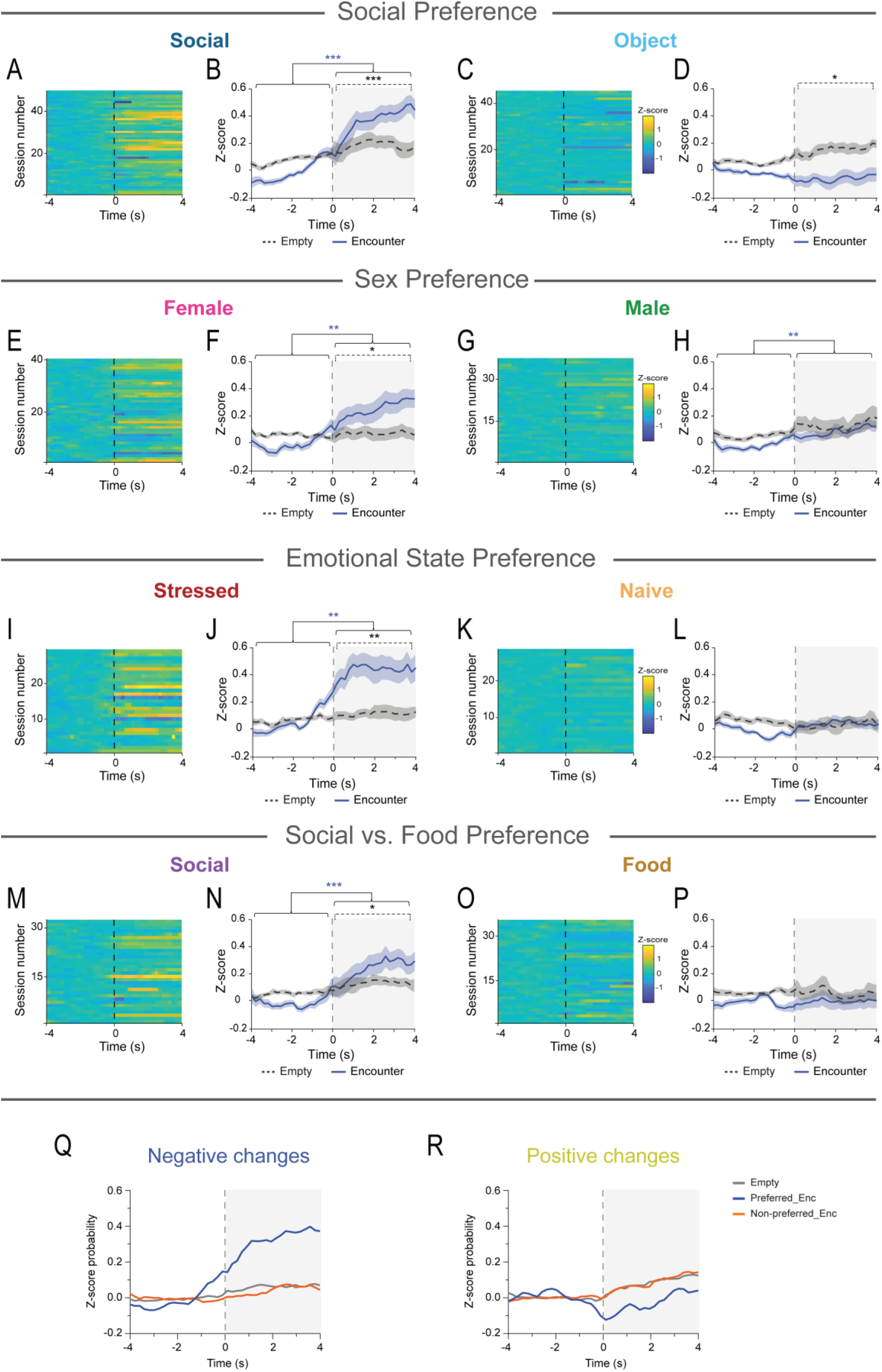
Negative calcium transients in mPFC pyramidal neurons’ activity predict social investigation behavior. **A.** Heat-map (see color code to the right of **C**) of the z-scored social stimulus investigation probability following (4 seconds in 0.2s bins) the onset of negative change events in the calcium signal (time ‘0’), averaged across all negative events within each session, with each row representing one session of the SP task. **B.** Superimposed traces of the mean (±SEM) social stimulus investigation probability for 4 seconds following the onset (time ‘0’) of negative change events identified in the pre-encounter stage with empty chambers (black dashed line) and in the encounter stage (dark blue filled line), z-scored to the probability in the (-4) - 0s window in 0.2 s bins, and averaged across all SP sessions. **C-D.** As in **A-B**, for the object in SP sessions. **E-H.** As in **A-D**, for the SxP task. **I-L.** As in **A-D**, for the ESPs task. **M-P.** As in **A-D**, for the SvFP task. **Q.** Mean traces for the investigation probability of the preferred stimulus (in blue) and the non-preferred stimulus (in orange) following negative change events in the calcium signal during the encounter stage of all sessions of the four tasks. Gray trace represents the mean investigation probability of empty chambers in the pre-encounter stage, later used to house the preferred and non-preferred stimuli. **R.** As in **Q**, for positive change events. Please note the high probability of investigating the preferred stimulus in the encounter stage compared to the other traces. *\*p<0.05*, \*\**p*<0.01, \*\*\**p*<0.001, One-sample Wilcoxon signed rank test (within trace differences in time) or Wilcoxon signed rank test (differences between pre-encounter and encounter).

Together, these findings suggest that decreased activity of mPFC pyramidal neurons serves a functional role in regulating investigative behavior toward the preferred stimulus and can be predictive of the animal’s stimulus investigation behavior.

### IV. Optogenetic stimulation of mPFC pyramidal neurons during stimulus investigation modulates social choice

To further examine whether and how mPFC pyramidal neurons play a critical role in mediating social choice, we used optogenetic stimulation to activate these neurons during stimulus investigation across all four tasks. For that, we used AAV viral vectors to express the excitatory opsin Channelrhodopsin-2 (ChR2) under the control of the CaMKII *α* promoter in the mPFC, followed by optic fiber implantation for blue light delivery (Fig. 5A). Histological analysis of the implanter optic fibers depth revealed that 6 out of 14 animals (42.85%) had the optic fiber tip located in the PrL, while 8 animals (57.15%) in the IL (Fig. S8A-B). Photo-stimulation (7–10 ms pulses at 10 Hz) was delivered specifically during investigation bouts toward a given stimulus (from onset to end of bout, Fig. 5B). Each task was repeated three times with the same animals under three distinct stimulation protocols (in random order): no stimulation, stimulation during bouts with one stimulus, and stimulation during bouts with the other stimulus. Notably, there were no differences in the total investigation time between the four tasks in the “no stimulation” condition, indicating a similar level of social motivation without stimulation (Fig. S8C).

**Figure 5.**
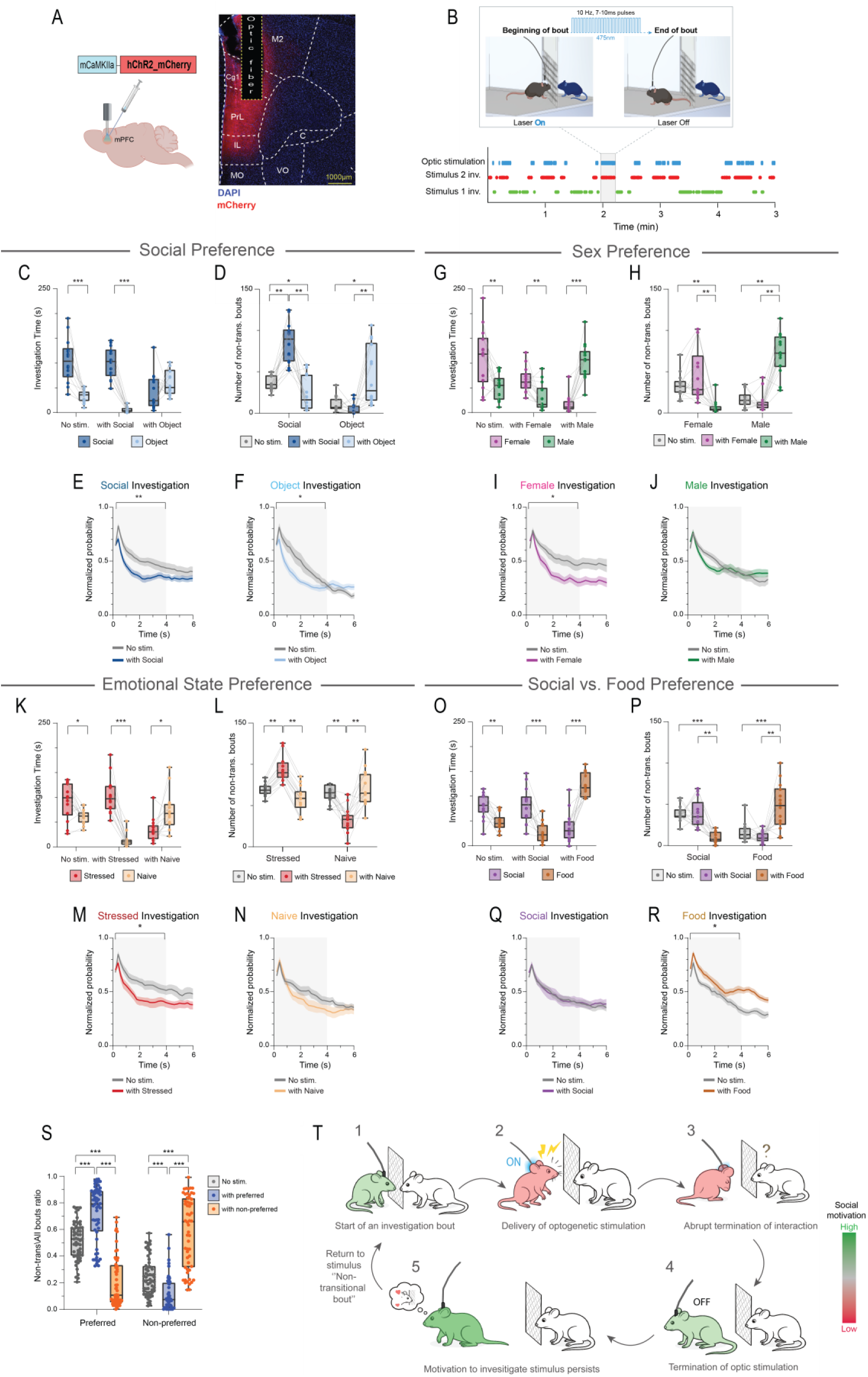
Optogenetic stimulation of mPFC pyramidal neurons during stimulus investigation modulates social choice. **A.** On the left, a schematic representation of the ChR2-expressing AAV vector injection followed by optic fiber implantation in the mPFC. On the right, a fluorescent micrograph of an infected mPFC slice, showing ChR2-mCherry expression (red) and the position of the optic fiber (dashed line). **B.** A session example for the investigation times of stimulus 1 (in green), stimulus 2 (in red), and periods of optic stimulation delivered during investigation bouts with stimulus 2 (in blue), across the 5 min duration of the session. The highlighted gray square represents an example bout with stimulus 2 in which optic stimulation (10ms pulses at 10Hz frequency) was delivered at the onset of the bout (when the subject’s nose touches the mesh of the stimulus chamber) and terminated at the end of the bout (when the subject turns away from the stimulus chamber) **C.** Median (in box plot^&^) investigation time of the social (dark blue) and object (light blue) stimuli (n=14), in SP sessions with no stimulation, or when optogenetic stimulation was consistently applied during investigation bouts with the social stimulus, or during investigation bouts with the object. A significant effect was found for stimulus (*p<0.001*), condition (*p<0.05*), and stimulus x condition interaction (*p<0.001*) by two-way RM ANOVA. **D.** Median (in box plot^&^) sum of non-transitional bouts to the social stimulus and the object across the three optic stimulation conditions of the SP task. **E.** Superimposed traces of the normalized mean (±SEM) probability for investigating the social stimulus 6 seconds, in 0.2s bins, following the bout onset in SP sessions with no optic stimulation (in gray), and in SP sessions with optic stimulation delivered during bouts with the social stimulus (in dark blue). The gray-shaded area represents the time window (0 to 4s) taken for statistical analysis. **F.** As in **E**, for the object stimulus in SP sessions with no optic stimulation (in gray), and in SP sessions with optic stimulation delivered during bouts with the object (in light blue). **G-J.** As in **C-F**, for the SxP task (n=14). **K-N.** As in **C-F**, for the ESPs task (n=14). **O-R.** As in **C-F**, for the SvFP task (n=14). **S.** Median (in box plot^&^) ratio of non-transitional to all bouts for the preferred and the non-preferred stimuli from all four tasks across the three optic stimulation conditions (no stimulation, stimulation with the preferred stimulus, and stimulation with the non-preferred stimulus). Please note the significant increase in the non-transitional/all bouts ratio for the stimulus coupled with the optic stimulation. A main effect for condition was found in Independent-Samples Kruskal-Wallis test for both preferred (*p<0.001*) and non-preferred (*p<0.001*) stimuli. **T.** Schematic representation of the behavioral sequence observed during optogenetic stimulation of pyramidal neurons in the mPFC. Subject mouse color indicates its social motivation level according to the color gradient on the right of the cartoon. *\*p<0.05*, \*\**p*<0.01, \*\*\**p*<0.001, *post-hoc* paired t-test with Holm-Šídák correction following detection of main effects by RM ANOVA in **G**, **K**, and **O**. *\*p<0.05*, \*\**p*<0.01, \*\*\**p*<0.001, Wilcoxon signed rank test with Holm-Šídák correction for multiple comparisons in **D**, **H**, **L**, and **P**. *\*p<0.05*, \*\**p*<0.01, independent t-test in **E-F**, **I-J, M-N,** and **Q-R**. *\*p<0.05*, \*\**p*<0.01, \*\*\**p*<0.001, *post-hoc* pairwise comparisons with Holm-Šídák correction for multiple comparisons following detection of main effects by Kruskal-Wallis test in **S**. ^&^ Box plot represents 25 to 75 percentiles of the distribution, while the bold line is the median of the distribution. Whiskers represent the smallest and largest values in the distribution.

We found that optogenetic stimulation delivered during investigation bouts with the preferred stimulus strengthened the preference toward it in all tasks except SxP, as reflected by the investigation time (Fig. 5C,G,K,O) and RDI values for the typically preferred stimulus (Fig. S8D-G). Conversely, stimulating mPFC pyramidal neurons during investigation of the non-preferred stimulus reverted the normal preference by creating a preference toward the typically non-preferred stimulus in the SxP, ESPs, and SvFP tasks as shown in both total investigation time (Fig. 5G,K,O) and RDI for the typically preferred stimulus. In the SP task, inverted preference to the object stimulus was found in the social stimulus RDI only (Fig. 5C; Fig. S8D). Thus, stimulating mPFC pyramidal neurons during stimulus investigation enhanced the preference toward that stimulus in all cases.

We then examined how optic stimulation of mPFC pyramidal neurons affects the structure of stimulus investigation. We first examined the number of transitional and non-transitional bouts toward each stimulus in all three stimulation conditions. In most cases we found a significantly greater number of non-transitional bouts toward the stimulus paired with optic stimulation (Supp. Movie 1-2), as compared to no stimulation, while the transitional bouts were affected less consistently (Fig. 5D,H,L,P; Fig. S8H-K). Accordingly, the ratio of non-transitional bouts to all bouts pulled across all tasks was significantly higher for the stimulation-paired stimulus compared to the other stimulus or non-stimulation condition, regardless of whether that stimulus is typically the preferred or the non-preferred (Fig. 5S).

These results suggest that optogenetic stimulation of mPFC pyramidal neurons elicits a high level of repetitive investigation bouts toward the stimulation-paired stimulus. In contrast, when analyzing the time course of all investigation bouts toward a given stimulus with and without stimulation, we found that in almost all cases (besides SvFP) optogenetic stimulation shortened the mean investigation bout rather than lengthening it (Fig. 5E-F,I-J,M-N,Q-R).

Thus, stimulation of mPFC pyramidal neurons during stimulus investigation exerts a mixed effect on investigation behavior, regardless of stimulus identity. On one hand, it shortened investigation bouts due to an immediate aversive effect that prompted the subjects to abruptly terminate their ongoing interaction with the stimulus. On the other hand, the motivation driving the subject to investigate the stimulus seems to persist despite the optogenetic stimulation. Therefore, once the optic stimulation is terminated, subject animals were more likely to resume investigating the same stimulus (i.e., perform a non-transitional bout). This led to an increased number of non-transitional investigation bouts, resulting in a counter-intuitive net increase in total investigation time for the stimulated stimulus (see cartoon in Fig. 5T).

### VI. mPFC pyramidal neurons generate an excitatory caution signal toward aversively conditioned social stimuli

Following the experiments described above, we hypothesized that excitatory activity of mPFC pyramidal neurons induces a ‘caution signal’ that promotes social avoidance. To directly test this hypothesis, we applied a social fear conditioning (SFC) paradigm (Fig. 6A), designed to invert the innately positive valence of a specific social stimulus into a negative valence by pairing its investigation with a foot shock. We recorded calcium signals from mPFC pyramidal neurons throughout all stages of the paradigm (pre-conditioning, early recall, and conditioned discrimination test) to track the neural correlates of the induced preference shift.

**Figure 6.**
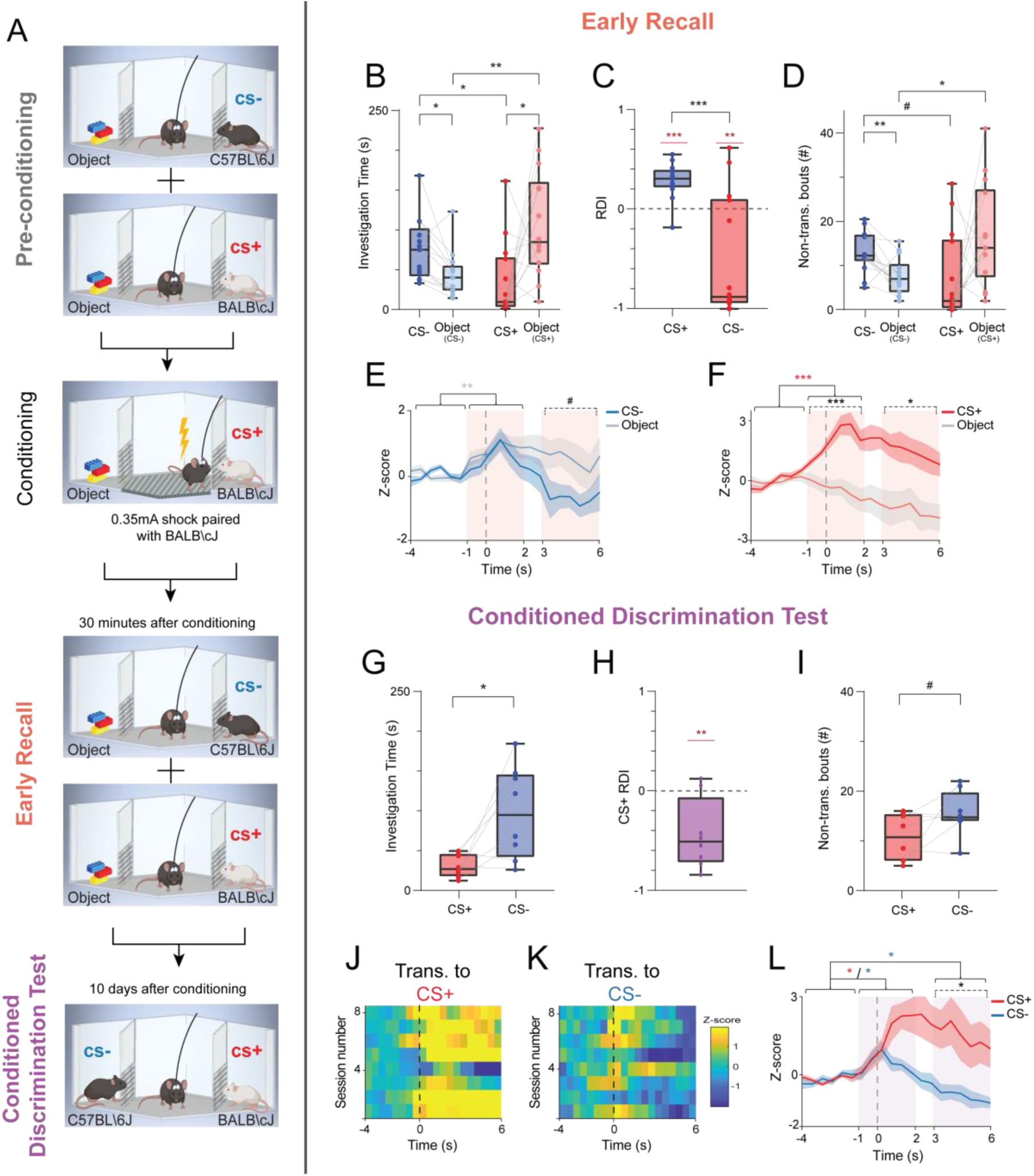
mPFC pyramidal neurons generate an excitatory caution signal toward aversively conditioned social stimuli. **A.** Schematic representation of the SFC paradigm. **B.** Median (in box plot^&^) value of mean investigation time of each of the social (C57BL/6J (CS-) and BALB/cJ (CS+)) and object stimuli in the 5-min early recall sessions (n=14 from 14 animals). A significant effect was found for stimulus x strain interaction (*p<0.01)* by two-way MM ANOVA. **C.** Median (in box plot^&^) relative discrimination index (RDI) for the social stimuli in the early recall sessions with CS+ and CS-. **D.** Median (in box plot^&^) value for the number of non-transitional bouts to the social (CS+ and CS-) and object stimuli in the early recall sessions. A significant effect was found for stimulus x strain interaction (*p<0.01)* by two-way MM ANOVA. **E.** Superimposed traces of the mean (±SEM) z-scored calcium signals averaged across all early recall sessions with CS- and object stimuli. Pink shaded areas represent two distinct time windows of the event which are used for the statistical Analysis. A significant effect was found for time (*p<0.05)* by two-way MM ANOVA. **F.** As in **E,** for the CS+ and object stimuli in the early recall sessions. A significant effect was found for stimulus *(p<0.01),* and time x stimulus interaction *(p<0.05)* by two-way MM ANOVA. **G.** Median (in box plot^&^) value of mean investigation time of CS- and CS+ in the 5-min conditioned discrimination sessions (n=8 from 8 animals). **H.** Median (in box plot^&^) relative discrimination index (RDI) for the CS+ in the conditioned discrimination sessions. **I.** Median (in box plot^&^) value for the number of non-transitional bouts to CS- and CS+ stimuli in the conditioned discrimination sessions. **J.** Heat-map (see color code to the right of **K**) of the z-scored calcium signals at the beginning (first six seconds) of bout, averaged across all transitional bouts toward the CS+ stimulus animals in the conditioned discrimination test. **K.** As in **J,** for the CS- stimulus. **L.** As in **E**, for the CS- and CS+ across all sessions of the conditioned discrimination test. A significant effect was found for time (*p<0.01),* stimulus *(p<0.05),* and time x stimulus interaction *(p<0.01)* by two-way MM ANOVA. *#p<0.07,* \**p<0.05, **p<0.01, post-hoc* independent t-test (for comparisons between stimuli from different tests) or paired t-test (for comparisons between stimuli from the same test) with Holm-Šídák correction for multiple comparisons following detection of main effects by ANOVA in **B**, and **D.** \*\**p*<0.01, \*\*\**p*<0.001 (in red), one-sample t-test with Holm-Šídák correction for multiple comparisons when present in **C,** and **H**. \*\*\**p*<0.001 (in black), independent samples t-test in **C**; *#p<0.07,* Paired t-test in **I**. *#p<0.07, *p<0.05,* \*\**p*<0.01, \*\*\**p*<0.001, *post-hoc* paired (within stimulus differences in time) or independent (differences between stimuli within time windows) t-test with Holm-Šídák correction for multiple comparisons following detection of main effects by ANOVA in **E-F**, and **L**. ^&^ Box plot represents 25 to 75 percentiles of the distribution, while the bold line is the median of the distribution. Whiskers represent the smallest and largest values in the distribution.

In the pre-conditioning stage, each subject mouse performed two SP tests, each with a social stimulus of a distinct strain, to make these stimuli easily distinguishable. Thus, in the first SP test, we used a C57BL\6J stimulus (CS-) and in the second test a BALB\cJ stimulus (CS+). The baseline tests were followed by a social fear conditioning (SFC) epoch, during which any approach by the subject to the BALB\cJ stimulus was paired with a mild foot shock (0.4 mA, typically 2–4 shocks). Then, two early recall SP tests, one with CS− and one with CS+ were conducted 30 min after the SFC epoch. Finally, for a direct comparison between the CS+ and CS- social stimuli, we conducted a conditioned discrimination test, where the fear-conditioned subject animals were exposed to both social stimuli in the same arena, ten days after the conditioning session.

First, we confirmed that the conditioning procedure successfully altered social preference, as we previously described [26]. During pre-conditioning, subject mice showed a clear preference for both CS- and CS+ social stimuli over an inanimate object. This preference was evidenced by the significantly longer time they dedicated to investigating the social stimuli over the objects (Fig. S9A), positive social stimulus RDI (Fig. S9B), and a significantly higher number of non-transitional bouts (Fig. S9C). In contrast, in the early recall phase that followed the SFC procedure, this preference was selectively altered. While subjects continued to prefer the CS- mouse, they now avoided the CS+ animal, which had been paired with the foot shock. This selective aversion was reflected by a significantly lower CS+ investigation time as compared to object investigation, a negative social stimulus RDI for the CS+, and no difference in the number of non-transitional bouts for CS+ compared to object (Fig. 6B-D). Next, we analyzed the corresponding changes in mPFC calcium signals, starting with the neural responses to empty chambers in the pre-encounter stage. While non-transitional bouts to empty chambers elicited no change in neural responses in both pre-conditioning and early recall sessions (Fig. S9F), transitional bouts exhibited increased neural activity in both stages. Yet, in early recall sessions, the excitatory response to empty chambers was significantly higher compared to pre-conditioning sessions (Fig. S9G). This result suggests that the conditioning procedure elevated the animal’s overall alertness and cautionary state, thereby amplifying the baseline excitatory signal associated with exploratory transitions.

We then examined changes in the mPFC calcium signal during the encounter stage, focusing on transitional bouts. In pre-conditioning, transitional bouts toward either the CS-or CS+ stimulus were associated with a slight, on-average, reduced mPFC pyramidal cell activity compared to the object stimulus (Fig. S9D-E), consistent with our earlier findings for preferred stimuli. Following conditioning, the neural response to the CS- retained the same pattern, showing the characteristic inhibition during transitions compared to the object (Fig. 6E). In contrast, the neural signature during transitional bouts toward the CS+ inverted markedly, now eliciting a significant excitatory response (Fig. 6F). Accordingly, a direct comparison of the responses to the CS+ stimulus before and after conditioning revealed a significant difference between them, with robust excitation in early recall compared to slight inhibition at pre-conditioning (Fig. S9H). Interestingly, the response to the object stimulus presented with the CS+ showed an opposite shift, with clear inhibition in early recall compared to slight excitation at pre-conditioning, reflecting the difference in the object’s relative value across the two stages of the paradigm (Fig. S9I). As for the object stimulus presented with the CS-, a different trend was observed, with higher responses in the early recall session than at pre-conditioning, even though the object remained the unpreferred stimulus in both stages of the SFC paradigm, while responses to the CS- remained comparable between the different stages of the SFC paradigm (Fig. S9J-K),

As expected, in the conditioned discrimination test, where the fear-conditioned subject animals were exposed to both the CS- and CS+ in the same arena, subject animals preferred the CS- animal over the CS+ animal (Fig. 6G-I). When mPFC calcium signals were examined, the CS+ elicited a robust excitatory response, while the CS- elicited the expected inhibition (Fig. 6J-L).

Taken together with our optogenetic stimulation results, the SFC results provide strong causal evidence for our ‘caution signal’ hypothesis. By transforming a preferred social stimulus into an aversive one, we induced a corresponding change in the mPFC transition-related signal, turning it from inhibition to a robust excitation. This demonstrates that excitatory activity in mPFC pyramidal neurons tracks the learned aversive value of a social target, functioning as a caution signal that drives subsequent social avoidant behavior.

### VII. Functional dissociation of mPFC projections’ role in mediating social choice

So far, we have examined the activity of the whole population of mPFC pyramidal neurons, assuming that they all function under a uniform activity organizing principle. However, sub-populations of these neurons project to several distinct brain regions [13], hence may have different principles organizing their activity during a binary choice between stimuli. To check this possibility, we examined the activity of two such sub-populations of mPFC pyramidal neurons: those that project to the nucleus accumbens (NAc) and those that project to the basolateral amygdala (BLA). To target mPFC neurons projecting to a given region we injected a retrogade CAV-Cre viral vector mixed with an AAV vector expressing mCherry (to label the injection site) into either the NAc or the BLA. We also injected into the mPFC an AAV vector leading to Cre-dependent expression of GCaMP7 under the human EF1α promoter, along with an optic fiber implantation (Fig. 7A-B). Upon recovery, we used fiber photometry to record calcium signals while subject animals performed the four tasks, as described above.

**Figure 7.**
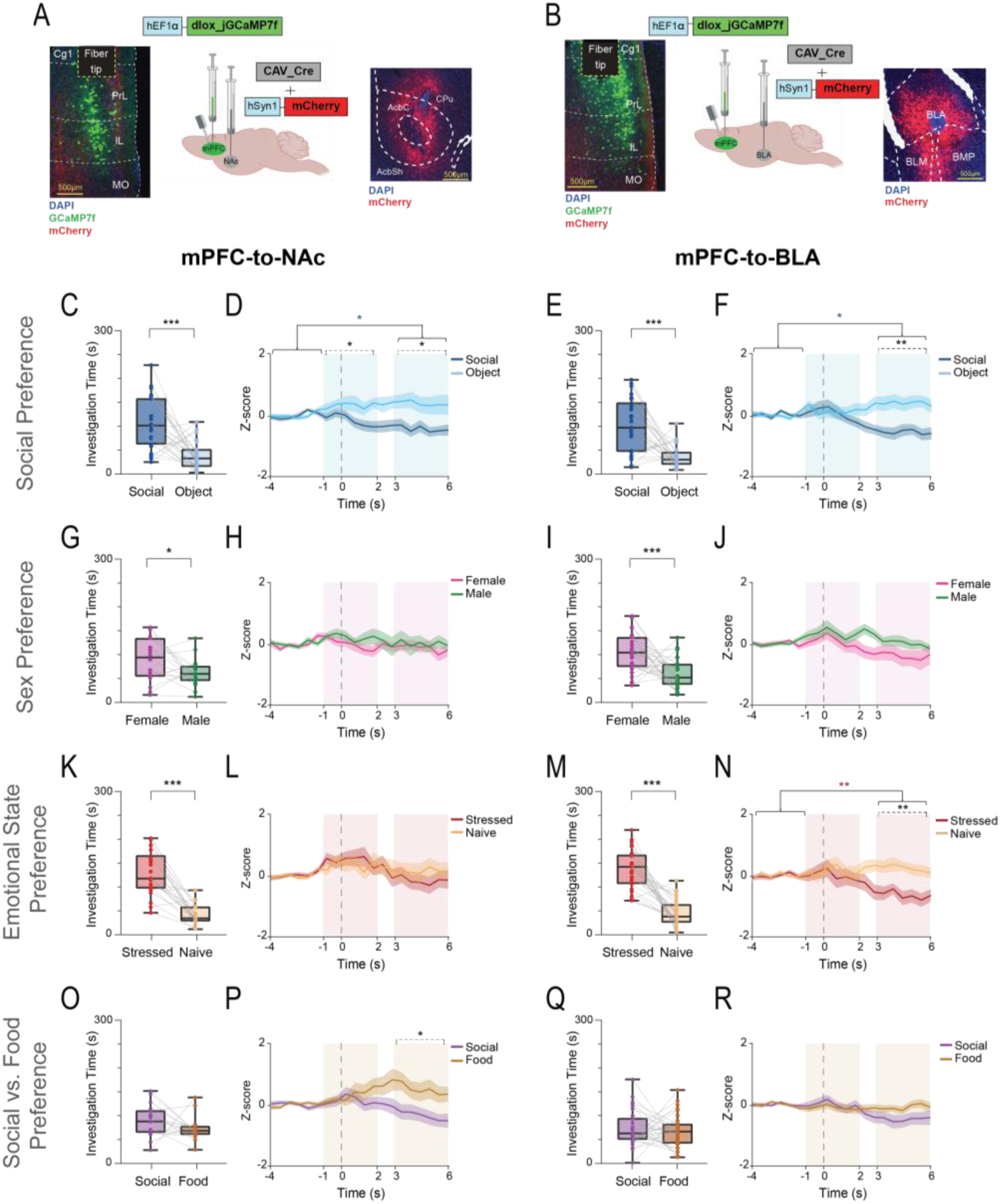
Projection-specific calcium dynamics in mPFC-to-NAc and mPFC-to-BLA neurons during transitional bouts. **A.** A schematic representation of the CAV_Cre and mCherry expressing AAV vectors injection into the NAc, and the dlox_GCaMP7f AAV vector injection into the mPFC, followed by optic fiber implantation in the mPFC. On the left, a fluorescent micrograph of an infected mPFC slice, showing GCaMP7f (green) and the position of the optic fiber (dashed line). On the right, a fluorescent micrograph of an infected NAc slice, showing mCherry (red) expression. **B.** On the left, as in **A**, for the injection and expression of GCaMP7f in mPFC-to-BLA projecting neurons. On the right, a fluorescent micrograph of an infected BLA slice, showing mCherry (red) expression. **C.** Median (in box plot^&^) investigation time of the social stimulus and the object (n=27 sessions from 11 animals) during the 5 min encounter stage of SP sessions recorded from animals with GCaMP7f expression in mPFC-to-NAc neuronal cells. **D.** Superimposed traces of the mean (±SEM) z-scored calcium signals for transitional bouts with the social stimulus (dark blue) and object (light blue), averaged across all SP sessions, and aligned to the bout onset (time ‘0’). Blue-shaded areas represent two distinct time windows of the event, which were used for the statistical Analysis. A significant effect was found for stimulus (*p<0.01*) and time x stimulus interaction (*p<0.01*) by two-way MM ANOVA. **E-F.** As **B-D**, for SP sessions recorded from animals with GCaMP7f expression in mPFC-to-BLA neuronal cells (n=25 sessions from 12 animals). **G-J.** As in **C-F**, for the SxP task (n_mPFC-to-NAc_=19 from 11 animals; n_mPFC-to-BLA_=31 from 12 animals). **K-N.** As in **C-F**, for the ESPs task (n_mPFC-to-NAc_=22 from 11 animals; n_mPFC-to-BLA_=27 from 12 animals). **O-R.** As in **C-F**, for the SvFP task (n_mPFC-to-NAc_=18 from 11 animals; n_mPFC-to-BLA_=32 from 12 animals). \**p<0.05*, *\*\*\*p<0.001*, paired t-test in **C**, **E**, **G**, **I**, **K**, and **M**. *\*p<0.05*, \*\**p*<0.01, *post-hoc* paired (within stimulus differences in time) or independent (differences between stimuli within time windows) t-test with Holm-Šídák correction for multiple comparisons following detection of main effects by ANOVA in **D,F,H,J,L,N,P**, and **R**. ^&^ Box plot represents 25 to 75 percentiles of the distribution, while the bold line is the median of the distribution. Whiskers represent the smallest and largest values in the distribution.

We first examined changes in mPFC calcium signal during non-transitional and transitional bouts to empty chambers in the pre-encounter stage (pooled from all recorded sessions), comparing responses across the three recorded neural populations: mPFC CaMKII*α*+ neurons, mPFC-to-NAc projecting neurons, and mPFC-to-BLA projecting neurons. In contrast to excitatory responses observed in the general population of pyramidal neurons during both bout types, neither the mPFC-to-NAc nor the mPFC-to-BLA projecting neurons exhibited comparable elevations in calcium signal (Fig. S10A-B). This dissociation indicates that, even in the absence of social stimuli, the two projection-defined subpopulations display distinct response profiles relative to the general population of pyramidal neurons, suggesting that their functional organization diverges from that of the general population already at the level of non-social, exploratory behavior.

As for encounter phase, in the SP, SxP, and ESPs tasks, subject animals of both groups (mPFC-to-NAc and mPFC-to-BLA) displayed a similar preference as reported earlier for the mCaMKII*α*-GCaMP6-injected animals (Fig. 7C,E,G,I,K,M), with the exception of the SvFP task, where subject animals from both groups displayed no preference for either stimulus (Fig. 7O,Q).

We next examined the neural activity of mPFC-to-NAc and mPFC-to-BLA projecting neurons during non-transitional and transitional bouts in each task. Similar to the general population of pyramidal neurons, differences in neural responses -when present- were only found during transitional bouts (Fig. 7C-R; Fig. S10C-R). Yet, these differential responses were task-dependent. In the SP task, both populations of projecting neurons exhibited a similar pattern of activity changes during transitional bouts, with inhibitory responses to the social stimulus specifically in transitional bouts (Fig. 7D,F). In the SxP task, neither population showed any significant response during transitional bouts (Fig. 7H,J). In ESPs, only mPFC-to-BLA projecting neurons showed inhibitory response during transitional bouts with the stressed stimulus (Fig. 7L,N). Conversely, In the SvFP task only the mPFC-to-NAc projecting neurons population exhibited inhibitory responses during transitional bouts with the social stimulus compared to the food (Fig. 7P,R).

Taken together, the mPFC-to-NAc and the mPFC-to-BLA projecting neurons exhibit distinct pattern of responses during social choice in a task-dependent manner. However, although their engagement varies across tasks, whenever these projections are recruited, their response profile follows the same consistent pattern observed in the broader population of mPFC pyramidal neurons: inhibition during transitional bouts toward the preferred stimulus.

## Discussion

Social decision-making requires the individual to continuously evaluate competing options and select appropriate actions. Here, we provide compelling evidence that the mPFC plays a central role in this process by encoding the context-dependent relative value of a social choice. Our principal finding is that the calcium signal recorded from mPFC pyramidal neurons exhibit selective inhibition during transitional bouts toward a preferred stimulus, and this inhibition predicts subsequent investigation behavior. This selective inhibition, which is contrasted with non-specific slight excitation taking place during the investigation of non-preferred stimuli, reveals a neural mechanism for social choice that is dynamically updated based on the relative appeal of available options.

Our results clearly demonstrate that the mPFC encodes the relative value of a social stimulus within a given context, rather than its absolute identity. This is exemplified by the “flipping” of the neural response to the same naïve male stimulus, which elicited inhibition when it was the preferred option (in the SP and SvFP tasks) as opposed to excitation when being non-preferred (in the SxP and ESPs tasks). This context-dependent encoding was further confirmed by the SvFP task, where social and food stimuli elicited opposite responses that flipped between satiety and food deprivation conditions, according to their relative preference. Taken together, our findings indicate that in the context of social choice, the mPFC employs a flexible but reliable, context-dependent coding scheme for stimulus value, implemented through the inhibition of pyramidal neurons. This mechanism integrates internal motivational states with external contexts to guide adaptive social choices, a flexibility that allows the same neurons to encode opposite values depending on the animal’s shifting priorities. Our findings are consistent with established principles of prefrontal cortex function. For instance, Hyman et al. (2012), showed that mPFC population activity is governed by the current task rules rather than by the physical attributes of stimuli [27]. Similarly, Walton et al. (2010) demonstrated that PFC responses to a given stimulus are continuously updated by context and past experience [28].

Additionally, we found that differential neural responses were primarily observed during transitional investigation bouts -when the subject animal switched between stimuli- and not during non-transitional bouts, which reflect repeated engagement with the same stimulus. These findings indicate that the mPFC is particularly involved in behavioral switching and the re-evaluation of choices, a core component of cognitive flexibility [29, 30], rather than in maintaining ongoing behavior. The inhibition observed during transitional bouts toward the preferred stimulus may reflect a neural “reset” that facilitates disengagement from the previous (non-preferred) option and commitment to the new (preferred) one. The lack of differential activity during non-transitional bouts may indicate that once a choice is made and the animal is engaged in sustained investigation, the mPFC’s role in value encoding becomes less critical, and other brain regions may take over the maintenance of the ongoing behavior. Recent work by Gabriel et al. (2025) provides important support for the selective engagement of mPFC in transitional bouts observed by us. These authors demonstrated that the PrL undergoes rapid reorganization of its population activity patterns during behavioral transitions in threat-avoidance learning. Critically, they showed that these reorganized neural representations emerge specifically at decision points where animals must discriminate between action outcomes and predict future choices, suggesting that the temporal dynamics of mPFC activity and its selective engagement during transitions may be a fundamental principle of prefrontal computation across diverse behavioral contexts [31].

While prior work has established a value-based role for PFC activity in guiding choice across non-social [32–36] and social [17, 20, 37–39] contexts, these findings predominantly derive from operant tasks with explicit rewards. In such tasks, animals learn to associate specific stimuli with reward outcomes through training, which may engage mPFC mechanisms primarily involved in executing learned associations. In contrast, our paradigm captures mPFC involvement in spontaneous social discrimination, where animals continuously evaluate social options based on their innate tendencies, without explicit conditioning. Such distinction is important because studies reporting predominantly excitatory mPFC responses during decision-making have typically employed operant or learned-behavior paradigms [7, 40, 41], whereas our inhibitory findings emerge from unconditioned, naturalistic social interactions. These differences suggest that the neural mechanisms underlying value encoding in the mPFC may differ fundamentally between spontaneous and conditioned social behaviors. In addition, while our fiber photometry recordings revealed a strong correlation between mPFC inhibition and social choice, this technique measures bulk calcium transients and may not directly reflect spiking activity. Moreover, it captures both somatic action potentials and dendritic inputs, with temporal dynamics slower than electrophysiological recordings [42, 43]. Recent work has demonstrated that fiber photometry signals correlate more strongly with neuropil calcium than with somatic calcium, suggesting they may reflect subthreshold changes in dendritic and axonal compartments rather than pure spiking activity [44].

Our optogenetic experiments provide causal evidence for the role of mPFC activity in regulating social choice. Stimulating mPFC pyramidal neurons during investigation of a given stimulus enhanced preference for that stimulus, regardless of its initial value. However, this preference enhancement occurred despite an immediate avoidance effect: stimulation shortened investigation bout duration and caused the animals to cease stimulus investigation. We propose that pronounced excitation in mPFC pyramidal neurons serves as a “caution” signal reflecting heightened vigilance or alertness. When this signal exceeds a certain threshold, it prompts disengagement from the investigated stimulus, explaining the immediate disruptive effects of optogenetic stimulation on stimulus investigation. Notably, multiple previous studies support this interpretation, showing that optogenetic stimulation of mPFC pyramidal neurons can produce negative or aversive effects on behavior, including enhanced fear associations and avoidance [45–50]. Upon stimulation termination, however, the animals rapidly and repeatedly re-engaged with the same stimulus, reflected by an increased number of non-transitional bouts. This pattern of repetitive returns resulted in a net increase in total investigation time and robust preference bias toward the stimulated option. The distinction between transitional and non-transitional bouts further supports this interpretation: we observed stronger excitation effects during transitional bouts between empty chambers compared to non-transitional bouts. Transitional bouts involve shifts between choices -moments of heightened uncertainty or caution as the animal evaluates a new option- whereas non-transitional bouts reflect sustained engagement with a familiar option. Thus, mild excitation during transitions to non-preferred options may reflect slight expression of the caution signal, promoting exploration of alternatives. Conversely, the inhibition exhibited by mPFC neurons during transitional bouts toward the preferred stimulus may reflect suppression of the caution signal, creating a “low-caution state” that facilitates engagement with that stimulus. This caution suppression is likely driven by inputs from limbic motivational circuits that process reward value and generate approach behavior (e.g. the basolateral amygdala (BLA), nucleus accumbens (NAc), and ventral tegmental area (VTA)) [14, 51–53]. Motivational signals from these limbic areas could preferentially activate local inhibitory interneurons (PV and SOM cells) [54, 55] in the mPFC, which then suppress pyramidal neuron activity to gate the caution signal [56].

Our social fear conditioning (SFC) results provide critical support for the caution signal interpretation and extend it to threat encoding. After a previously neutral social stimulus was paired with aversive foot shock, we observed a robust increase in mPFC pyramidal activity during transitional bouts toward that stimulus. This finding directly parallels recent work by Gu & Johansen (2025), who demonstrated that aversive conditioning induces heightened dmPFC excitability for both directly threat-paired and indirectly associated stimuli as part of an “internal model for emotional inference” [57]. Our findings, therefore, indicate that the mPFC generates context-dependent caution signals, implemented through pyramidal excitability, to encode learnt threat and promote cautious, avoidant responses. During social interactions, inhibition of this signal permits commitment to preferred options, while during threat encounters, expression of this signal promotes defensive avoidance. These caution signals are likely routed to downstream fear and anxiety circuits to guide adaptive defensive behaviors, representing a unified neural mechanism for flexible social decision-making across affiliative and aversive contexts.

The combined results from our social preference and fear conditioning experiments suggest that the mPFC operates on a “common currency” of value that encompasses both positive and negative social outcomes [58, 59]. In our initial experiments, the degree of pyramidal cell inhibition during transitional bouts correlated with the relative preference for a social partner, encoding a positive decision variable. Conversely, in the SFC experiment, the degree of excitation correlated with the learned aversion for a social partner, encoding a negative or threatening value. Together, these findings indicate that a given neural population can flexibly encode both the valence and magnitude of social value, spanning highly appetitive to highly aversive states. This framework is consistent with the common currency hypothesis, which posits that the mPFC integrates diverse inputs into a unified value signal to guide choice [58, 60]. The integration of internal state and external context into a unified value signal is further illustrated by the reversal of the mPFC response in the SvFP task following food deprivation. This suggests that internal state gates the sign of the population signal, consistent with models in which PFC integrates motivational and interoceptive information into valuation and choice [61].

The mPFC’s influence on social choice likely operates through multiple parallel output channels rather than a single centralized signal [6, 62]. Our recordings from mPFC neurons projecting to the NAc and BLA indicate that these projection-specific populations exhibit distinct functional specializations that depend on the behavioral choice context. Notably, this functional dissociation was evident even in non-social contexts: unlike the general CaMKII*α*^+^ pyramidal population, neither the mPFC-to-NAc nor the mPFC-to-BLA projecting neurons exhibited elevated excitatory responses during transitional or non-transitional bouts toward empty chambers in the pre-encounter stage. This suggests that the excitatory responses to empty chambers observed in the global population are not carried by these two populations, and that other populations of mPFC output neurons contribute to this signal. One candidate is the mPFC projection to the mediodorsal thalamus (MD), which has been implicated in task engagement and behavioral flexibility, with disruption of the mPFC-to-MD circuit impairing the ability to sustain attention and adapt behavior across changing contexts [29]. Another candidate is the mPFC projection to the dorsal striatum, which has been shown to mediate value-based choice through prelimbic circuits targeting distinct dorsal striatal subregions, and whose activity tracks expected outcome value during exploratory behavior [35, 36].

When social stimuli were present, mPFC-NAc and mPFC-BLA projecting neurons exhibited the same inhibitory response to preferred stimuli observed in the global population, but were engaged in a task-dependent manner. Our findings align with a growing body of literature demonstrating functional specialization of mPFC output pathways [13, 63, 64]. For instance, recent work has shown that mPFC-NAc projections more strongly track reward-related and social dominance signals [10, 63, 65–67], whereas mPFC-BLA projections are more involved in processing negative emotional valence and social novelty [45, 57, 64, 68, 69]. The distinct responses we observed in mPFC-NAc and mPFC-BLA projecting neurons may reflect a division of labor, where one pathway signals the value of the chosen option to reward-related circuits, while the other signals the emotional or social context to valence-processing regions. This projection-specific routing is also consistent with the common currency hypothesis, which predicts that value is computed centrally (in the mPFC) but distributed to multiple downstream systems for different functions [60]. Future studies using projection-specific manipulations, including recordings and targeted interventions in mPFC-to-thalamus and mPFC-to-striatum pathways, as well as selective manipulation of the mPFC-to-NAc and mPFC-to-BLA projections characterized here, will be necessary to fully dissect how social choice signals are routed and implemented across downstream circuits.

Our findings have several broader implications for understanding the neural basis of both typical and atypical social behavior, particularly concerning the balance of excitation and inhibition (E/I) in the mPFC. The strong, state-dependent inhibition we observed is likely implemented by specific interneuron subtypes that gate the expression of internal value models at the population level. This aligns with work showing that subtle shifts in the recruitment of inhibitory interneurons, such as parvalbumin (PV) [4, 70] or somatostatin (SST) [71] expressing cells, can qualitatively reshape how cortical populations represent social information and decision variables [4, 72–74]. The integrity of this inhibitory control is critical, as a disruption in E/I balance in the mPFC is a hallmark feature of Autism Spectrum Disorders (ASD) and has been proposed to underlie many of its core social deficits [73, 75, 76]. Our findings suggest a specific mechanism through which this may occur: if the inhibitory signal that marks commitment to a preferred social option is weakened or absent, individuals may have difficulty committing to social interactions or fail to appropriately prioritize social stimuli. This hypothesis is supported by studies demonstrating that restoring E/I balance in the mPFC can rescue social behavior deficits in mouse models of ASD [70, 72, 76]. Therefore, future studies examining whether the context-dependent value encoding mechanism we observed is disrupted in models of ASD will be important for understanding the neural basis of social deficits and for developing targeted interventions.

A key limitation of our study is that our recordings did not allow us to precisely target either the PrL or IL subregions of the mPFC, often suggested to play distinct or even opposing roles in behavior [15, 16, 77–79]. Our analysis revealed no significant correlations between optic fiber depth and neural activity during either transitional or non-transitional bouts, suggesting that the observed mPFC pyramidal cell responses are a general phenomenon across both subregions. This lack of subregional specificity is consistent with findings in other behavioral contexts where both PrL and IL have been shown to be similarly involved, such as in the processing of appetitive contextual memories and in the inhibitory control of reward-seeking behavior [80, 81]. However, it is possible that the inhibitory and excitatory signals we observed arise from different subregions or layers with varying weightings depending on the specific discrimination task, or that the relative contributions of PrL and IL vary in task contexts in ways not captured by our depth-based analysis. Future studies using more targeted approaches will be necessary to dissect the specific contributions of the PrL and IL to social choice and to determine whether they contribute differentially to the cautious signal hypothesis.

Second, our results are based on fiber photometry recordings which do not provide the single-cell resolution needed to understand the underlying circuit dynamics in detail [82]. Single-unit recordings or calcium imaging with cellular resolution will be necessary in future studies to elucidate the precise interplay of excitatory and inhibitory neurons that gives rise to the observed population-level signals. Interneurons may also be involved in the inhibitory signal we observed during interactions with preferred stimuli, perhaps by suppressing pyramidal neuronal activity when the animal commits to a preferred option. Future studies using cell-type-specific manipulations will be necessary to test this hypothesis.

In conclusion, our study provides novel insights into the neural mechanisms of social choice, demonstrating that the mPFC encodes the relative value of social options through a context-dependent inhibitory signal that is dynamically updated based on the subject’s behavioral state and motivational context. These findings not only advance our understanding of the social brain but also provide a new framework for investigating the neural basis of social deficits in neuropsychiatric disorders. By revealing that the mPFC operates on a common currency that integrates both positive and negative social values, and by identifying selective inhibition as a key mechanism for suppressing caution signals and enabling commitment to preferred social options, our work establishes fundamental principles of how the prefrontal cortex guides adaptive social decision-making.

## Supporting information

An Excel file containing details of all statistical analyses, organized by figure.

An Excel file containing processed datasets analyzed and organized by figure.

An Excel file containing optogenetic stimulation parameters and data retention rate.

Description of Additional Supplementary Files

## Declarations

## Acknowledgments

We thank Boris Shklyar, Head of the Bio-imaging Unit and Eng. Alex Bizer, the experimental systems engineer of the Faculty of Natural Sciences of the University of Haifa, for their technical help.

## Author contributions

R.J.: Formal Analysis, Investigation, Methodology, Validation, Visualization, Writing - original draft, and Writing - review & editing; P.J.: Investigation, Analysis, Methodology, and Visualization; S.N.: Data curation, Project administration, Software, Validation, Visualization, Writing - original draft, and Writing - review & editing. S.W.: Conceptualization, Funding acquisition, Project administration, Resources, Supervision, Writing - original draft, and Writing - review & editing.

## Additional Supplementary Files

**Table S1**. An Excel file containing details of all statistical analyses, organized by figure.

**Table S2**. An Excel file containing processed datasets analyzed and organized by figure.

**Tables S3**. Contains optogenetic stimulation parameters and data retention rate.

**Supplementary Movie 1**. Example clip (37s long) from an SP session with optic stimulation delivered during bouts with the social stimulus.

**Supplementary Movie 2**. Example (31s long) clip from an ESPs session with optic stimulation delivered during bouts with the stressed stimulus.

**Supplementary Movie 3**. Example (20s long) clip from a SxP session with optic stimulation delivered during bouts with the Female stimulus.

## Methods

### Method details

#### Resource availability

Further information and requests for resources and reagents should be directed to and will be fulfilled by Shlomo Wagner.

#### Materials availability

This study did not generate new unique reagents.

#### Data and code availability

All the data generated from experiments and the custom codes used to evaluate the conclusions of this paper would be uploaded onto a public repository upon acceptance of the paper.

### Experimental model and subject details

#### Animals

<u>C57BL/6J</u>: C57BL/6J naïve adult (8-18 week-old) male and female mice (Envigo RMS, Israel) were used either as subjects or stimuli. BALB/cJ naïve adult (8-18 week-old) males were used as stimuli in the Social Fear Conditioning paradigm.

<u>Maintenance:</u> Animals were housed in groups of 2-5 sex-matched mice per cage. All behavioral sessions were conducted while subjects were group-housed, with the exception of the SP task in CaMKII*α*–GCaMP6f–injected mice, which were recorded after two to three weeks of individual housing to enhance social motivation. All animals were housed at the animal facility of the University of Haifa under veterinary supervision, in a 12 h light/12 h dark cycle (lights on at 9 PM), with *ad libitum* access to food (standard chow diet, Envigo RMS, Israel) and water.

#### Institutional review board

All experiments were performed according to the National Institutes of Health guide for the care and use of laboratory animals and approved by the Institutional Animal Care and Use Committee of the University of Haifa.

#### Experiments

Behavioral experiments were conducted in the dark phase of the dark/light cycle in a sound attenuated chamber, under dim red light.

#### Experimental setups

The experimental setup [83] consisted of a white Plexiglas arena (37 X 22 X 35 cm) placed in the middle of an acoustic cabinet (60 X 65 X 80 cm). Two Plexiglas triangular chambers (12 cm isosceles, 35 cm height), into which an animal or object (plastic toy) stimulus could be introduced, were placed in two randomly selected opposite corners of the arena. A metal mesh (12 X 6 cm, 1 X 1 cm holes) placed at the bottom of the triangular chamber allowed direct interaction with the stimulus animals through the mesh. Food pellets, when used as stimulus, were placed in a chamber with a metal mesh with smaller holes (0.5 × 0.5 cm) that prevented subjects from grabbing or consuming the food during testing, thereby minimizing distraction and maintaining engagement with the experimental task. A high-quality monochromatic camera (Flea3 USB3, Flir for optogenetic experiments or Grasshopper3 for fiber photometry recordings) equipped with a wide-angle lens was placed at the top of the acoustic chamber and connected to a computer, enabling a clear view and recording (∼30 frames/s) of subject behavior using commercial software (FlyCapture2 or SpinView, FLIR).

#### Behavioral paradigms

All behavioral paradigms started with 15 minutes of habituation in which subject animals were moved into the experimental arena with two empty chambers. Throughout this time, social stimuli were placed in their chambers near the acoustic cabinet for acclimation. After habituation, the chambers containing the relevant stimuli were diagonally placed at opposite ends of the arena in a random fashion. Each task was conducted for 5 minutes, and differed mainly in the stimuli presented to the subjects-

Social preference (SP): Subject mice were introduced to either a social (naïve, novel, age-and sex-matched conspecific) and an object (mouse-size lego block).

Sex Preference (SxP): subject mice were presented with a novel male on one side, and a novel female conspecific on the other.

<u>Emotional State Preference for stress stimuli (ESPs)</u> - subject mice were introduced to a naive stimulus animal and to another stimulus animal that had been constrained for 15 min in a 50 ml tube pierced with multiple holes for ventilation. Each “stressed” stimulus animal was used as a stimulus for one session only.

<u>Social vs. food (SvFP, in satiety)</u>: Subject mice were introduced to a novel, naïve, social stimulus, and to food pellets composed of standard chow, under satiety conditions (*ad libitum* access to food and water prior to the task). Food pellets were placed inside a dedicated chamber fitted with a metal mesh with small holes (0.5 × 0.5 cm) that prevented subjects from grabbing or consuming the food during testing, thereby minimizing distraction and maintaining engagement with the experimental task.

<u>Social vs. food (SvFP, in food deprivation)</u>- subject mice were food deprived, with full access to water for 24 hours before the task. In the task, subjects were introduced to a novel, naïve, social stimulus, and to food pellets composed of standard chow.

<u>Social Fear Conditioning (SFC) Paradigm-</u> The Social Fear Conditioning task is composed of multiple stages. To establish innate social preference, subject mice were first moved to the testing arena with two empty chambers for 15-min habituation, followed by two consecutive SP tests with social stimuli of two distinct strains (C57BL/6J as CS-, and BALB/cJ as CS+), separated by a 15-min interval (this stage of the paradigm is termed “pre-conditioning”). Twenty minutes after the second SP test, subjects were transferred to the SFC arena. The arena was equipped with a removable stainless-steel grid floor (H10-11M, Coulbourn-Instruments) connected to a shock-delivery apparatus (precision regulated animal shocker H13-14, Coulbourn-Instruments), allowing manual application of foot shocks. Subjects underwent a 15-minute habituation period, followed by 5 minutes of the SFC procedure. During the SFC procedure, the subject received a mild electrical foot shock (0.3–0.4 mA, 750 ms duration) each time it initiated an investigation bout with the chamber containing the same BALB/cJ stimulus from baseline. Following conditioning, subjects were then moved to fresh cags where they were kept individually for 30 minutes. Following, two consecutive SP tests with the same stimuli as before conditioning were conducted (this phase of the paradigm is termed “Early Recall”). The animals were then placed back in their original home-cages, and 10 days later, they were returned to the experimental arena for 15-minute habituation followed by a 5-minute task in which subjects were presented simultaneously with the same CS- and the conditioned CS+ stimuli (Conditioned Discrimination Test, CDT). Fiber photometry recordings of mPFC pyramidal neurons were acquired throughout all sessions of the experiment.

#### Stereotactic surgery for viral injection and optic fiber implantation

<u>Surgery</u>: Mice were anesthetized and anesthetized with an intraperitoneal injection of a mixture of Ketamine, Domitor, and the painkiller Meloxicam (0.13 mg/g, 0.01 mg/g, and 0.01 mg/g, respectively). Anesthesia levels were monitored by testing toe pinch reflexes and maintained using isoflurane (0.2%; SomnoSuite, Kent Scientific Corporation). Body temperature was kept constant at approximately 37°C using a closed-loop custom-made temperature controller connected to a temperature probe and a heating pad placed under the animal. Ophthalmic ointment (Duratears, Alcon, Couvreur, NV) was applied to maintain eye moisture. Anesthetized animals were fixed in a stereotaxic apparatus (Kopf), with the head flat. The head was shaved, the scalp was disinfected, and the skin was removed to expose the skull. The skull was then leveled using bregma–lambda measurements.

The region of interest was then marked, and a hole (unilateral, right hemisphere) was drilled for viral injection and optic fiber implantation. Additional holes were drilled for supporting screws. Viral injection was performed as previously described in detail [26] using a glass capillary filled with the virus ssAAV-1/2-mCaMKII *α* -GCaMP6f-WPRE-SV40p(A) for fiber photometry recordings or ssAAV-1/2-mCaMKII *α* - hChR2(H134R)_mCherry-WPRE-SV40p(A) for excitatory optogenetic stimulation, was slowly lowered into the mPFC (AP, +2 mm; ML, -0.35 mm; DV, +2.0 mm) and left in place for 5 min prior to injection and 10 min following injection to prevent retraction of the virus. For targeting mPFC projecting neurons to the NAc and the BLA, CAV2-Cre (Canine Adenovirus Type 2 expressing Cre recombinase) in combination of 1:5 ratio with ssAAV-1/2-hSyn1-hChR2(H134R)_mCherry-WPRE-hGHp(A) for marking the injection site were injected into either the NAc (AP, +1.42 mm; ML, -1.2 mm; DV, +4.4 mm) or the BLA (AP, -1.43 mm; ML, -3.0 mm; DV, +4.8 mm). ssAAV-1/2-hEF1a-dlox-jGCaMP7f(rev)-dlox-WPRE-bGHp(A) was injected into the mPFC (AP, +2 mm; ML, -0.35 mm; DV, +2.0 mm). The Cav-cre virus was obtained from the Plateforme de Vectorologie de Montpellier (PVM), Institut de Génomique Fonctionnelle, CNRS, University of Montpellier, France. All other viruses were purchased from the Viral Vector Facility (VVF), Institute of Pharmacology & Toxicology, University of Zurich, Switzerland.

A total of 300 nl of the relevant virus was then delivered by manual application using a 50 ml syringe connected to the glass capillary through a plastic tube for applying air pressure (BRAND, disposable BLAUBRAND micropipettes, intraMark, 5 *μl*). Following viral injection, an optic fiber (Doric lenses, 200 μm, NA 0.66, 3 mm long, zirconia ferrule, flat-fiber tip) was inserted into the mPFC with the exact coordinates as the viral injection, and placed 200 *μ*m above the injection depth. Screws and optic fiber were fixed to the skull plate using dental cement (UNIFAST, GC America).

After the surgery, Antisedan (0.1 mL/10g body weight) was administered subcutaneously to wake the mice from anesthesia. The mice were injected with meloxicam (5%, 0.01 mg/g) and baytril (5%, 0.03 ml/10g) to relieve pain and prevent infections for three days following surgery. Behavioral testing with optogenetic stimulation or fiber photometry was conducted three weeks after surgery to allow optimal viral expression. Following surgery, all subjects were housed with one or two cage mates.

#### *In vivo* fiber photometry recordings

<u>Test</u>: All fiber photometry recordings were performed using the apparatus described above. The optic fiber of the implanted mouse was connected to the optical patch cords via a sleeve connector (Doric) under brief isoflurane anesthesia. The wired animal was then allowed to habituate in the behavior setup for 15 minutes. The recording session started with an additional 5-minute baseline (pre-encounter) period with empty chambers. Thereafter, the empty chambers were replaced with chambers containing the appropriate stimuli for a 5-minute test. Each subject was recorded in two sessions of distinct tasks per day, with at least 30 minutes between sessions for the same animal. Following the two sessions, animals were returned to their home-cage.

<u>Fiber photometry recording and synchronization to video recording</u>: Calcium signals were recorded using one of two fiber photometry recording setups composed of the RZ10x system of Tucker Davis Technologies and an optical path by Doric, which includes 600 μm mono-fiber optic patch cords connected to either a four ports Fluorescence MiniCube (FMC4_IE(400-410)_E(460-490)_F(500-550)_S) for one-color recordings or a six ports Fluorescence MiniCube (FMC6_IE(400-410)_E1(460-490)_F1(500-540)_E2(555-570)_F2(580-680)_S) for two-colors recordings. Optic paths were connected using a 200 μm optical patch cord with a fiber-optic rotary joint (FRJ_1×1_PT_200/230/LWMJ-0.57_1m_FCM_0.15m_FCM) that was connected to the recorded animal. A camera (Grasshopper3 USB3, FLIR) was placed on top of the arena. The Camera was connected to the digital port of the RZ10× system through its GPIO connector and configured to deliver strobes for every frame acquired at a frame rate of 30 frames/sec. These were later used to synchronize the video frames with the calcium signal. TDT Synapse software was used to record the GCaMP6f/7f signal channel (excitation at 465 nm, modulated at 330 Hz), the isosbestic control channel (405 nm, modulated at 210 Hz), and the digital channel receiving the camera strobes, with a sampling rate of ∼1017.3 Hz. LED currents were adjusted to return the voltage to between 0 and 200 mV for each signal, were offset by 5 mA, and were demodulated using a 6 Hz low-pass frequency filter. Light intensity levels for the 465-nm and 405-nm channels were verified by measuring output power with an external power meter (PM100D, Thorlabs) every 2–3 recording days. Output power was kept within the range of 11-18*μ*W for the 405-nm channel, and 20-30*μ*W for the 465-nm channel.

#### *In vivo* optogenetic stimulation

<u>Test:</u> In all optogenetic experiments (SP, SxP, ESPs, SvFP–Satiety, and SvFP–Food Deprivation), subject mice were connected to the stimulation apparatus and allowed to habituate for 20 minutes in an arena containing empty chambers. After habituation, the empty chambers were replaced with the appropriate stimuli chambers, and interactions were recorded for 5 minutes. In control experiments with empty chambers, new empty chambers were introduced into the arena at this stage.

Each subject was tested three times in the SP, SxP, ESPs, and SvFP tasks, each session employing a different stimulation condition: (1) no stimulation, (2) stimulation delivered during investigation bouts with one of the two presented stimuli, and (3) stimulation delivered during investigation bouts with the other stimulus. Optic stimulation during an investigation bout was manually delivered and defined as light delivery from the onset of nose contact with the mesh of a stimulus chamber until the subject’s head turned away from it.

Each day, subject mice participated in either a single experiment with optogenetic stimulation or in two sequential experiments, consisting of a non-stimulated session followed 20 minutes later by a stimulated session. All tasks were conducted in a randomized order.

<u>Optogenetic stimulation:</u> Subject mice were lightly anesthetized with isoflurane and connected to the laser (473 nm, model FTEC2471-M75YY0, Blue Sky Research) via a fiber-optic rotary joint attached to 200 µm core multimode fiber patch cables connecting the animal to the light source for optic stimulation delivery (Thorlabs, Fiber Optic Rotary Joint Patch Cables for Optogenetics, product group #6865). Animals were given 20 minutes to recover from any residual effects of the brief isoflurane anesthesia before testing, until they showed full mobility. In all experiments, optogenetic stimulation consisted of continuous light pulses delivered at 10 Hz throughout the entire duration of each investigation bout with a given stimulus. The initial photo-stimulation protocol used 10 ms pulses at a light power of 8.23 mW. However, these parameters induced seizure-like behavior in a subset of animals, which we attributed to the high level of ChR2 expression in the mPFC. This phenomenon has been previously reported in optogenetic studies of cortical regions, where brief optogenetic stimuli can elicit seizures that increase in severity with repeated stimulation (Cela et al., 2019; Osawa et al., 2013). To ensure that experimental outcomes reflected controlled neuronal modulation rather than pathological network states, we systematically optimized the stimulation parameters by first reducing light power from 8.23 mW to no lower than 4.65 mW, and subsequently shortening pulse duration from 10 ms to 7 ms, while maintaining the 10 Hz frequency (see Tables S3A-B for detailed timeline and voltage-light power calibration). The minimal used parameters of 4.65 mW and 7 ms pulses are well within the established range for effective ChR2 activation in vivo. Using a computational model based on direct measurements in mammalian brain tissue [84] (https://web.stanford.edu/group/dlab/cgi-bin/graph/chart.php), we calculated that 4.65mW of 473nm light delivered through a 200µm fiber (NA 0.66) produces an irradiance of approximately 4.99mW/mm² at 250µm distance from the fiber tip (the maximal distance between the fiber tip and the virus injection site), and 1.26mW/mm2 at a distance of 500µm which is the highest penetration depth of blue light stimulation in rodent brain tissues [49]. These values exceed the generally accepted threshold of 1 mW/mm² required for suprathreshold activation of ChR2-expressing neurons [84–86], while remaining within the recommended safe operating range of less than 10-20 mW/mm² at target cells to minimize tissue heating [49]. Furthermore, our pulse duration of 7 ms is well above the opening rate time constant of ChR2 (τ ≈ 1.21 ms), ensuring sufficient channel opening to trigger action potentials reliably [87]. Optical stimulation was manually delivered via a Master8 Channel Programmable Pulse Stimulator (A.M.P.I) connected to the laser. Sessions in which severe seizures prevented completion of behavioral testing were excluded from analysis. Overall, 12 out of 150 stimulation sessions (8%) were excluded due to severe seizures, yielding a 92% data retention rate across all animals (Table S3C). Importantly, even at the optimized parameters (4.65 mW, 7 ms pulses, 10 Hz), animals continued to exhibit the intended behavioral response pattern, characterized by abrupt termination of investigation bouts upon stimulation onset, followed by immediate re-engagement with the same stimulus once stimulation ceased, confirming that the optimized parameters were sufficient to produce the desired modulatory effect on mPFC pyramidal neuron activity (see Supp. Movie 3).

#### *Post-mortem* histological Analysis of viral injections and optic-fiber placement

Each implanted mouse was anesthetized with isoflurane (induction: 3%, 0.5%-0.8% maintenance in 200 mL/min of air), attached to the stereotaxic device, and the optic fiber implant was gently extracted from its head. The mouse was then perfused with phosphate-buffered saline (PBS) and fixed using a 4% paraformaldehyde (PFA, Sigma) solution. The brains were harvested and placed in PFA (4%) for 48 hours, followed by sectioning into 50-μm slices in the horizontal axis using a VT1200s Leica sliding vibratome. The slices were collected onto microscope slides and stained with DAPI (Fluoroshield with DAPI, histology mounting medium, Merck). Images of subject brain slices were acquired for verification of the viral expression placement and optic fiber path within the mPFC using an epifluorescence microscope (Ti2 eclipse, Nikon) equipped with a blue filter for DAPI staining, FITC for detecting GCaMP6f/7f-expressing cells, and TRITC for detecting mCherry-expressing cells.

### Quantification and statistical analysis

All processed data are detailed in Table S2, organized by the various figures.

#### Tracking software and behavioral analyses

All video recordings were analyzed using *TrackRodent* (https://github.com/shainetser/TrackRodent) [88]. The *BlackMouseWiredBodyBased* algorithm was applied to sessions involving tethered animals in both fiber photometry and optogenetic experiments. Behavioral analyses were performed as previously described [89].

#### Parameters for behavioral quantification

All behavioral parameters reported in this study were automatically extracted using DeepPhenotyping software [90, 91], which continuously tracks the animal’s position and investigation of stimuli chambers throughout the experiment. Behavioral events were identified and quantified using predefined algorithms and rule-based classifications embedded in the software. preference) minus the investigation time for the other stimulus, divided by the sum of both.

Investigation bouts were defined as events beginning when the animal’s head (or body center, depending on the selected algorithm in TrackRodent software) made contact with the metal mesh of the stimulus chamber and ending when the animal disengaged from the mesh. Each bout was subsequently categorized as either non-transitional or transitional. Non-transitional bouts were those preceded by a bout directed toward the same stimulus, whereas transitional bouts were those preceded by a bout directed toward the alternate stimulus. A minimum interval of >0.5 s between successive bouts was required to classify bouts as distinct. Behavioral events of interest detected by TrackRodent were then aligned with the calcium imaging data to enable precise synchronization between neural activity and behavior.

The Relative Discrimination Index (RDI) was calculated as the difference in investigation time between the two stimuli—specifically, investigation of the preferred stimulus minus the investigation of the non-preferred stimulus—normalized by the total investigation time toward both stimuli. For the naïve male stimulus (the common stimulus across all four tasks), the RDI (named “common stimulus discrimination index”) was calculated as the investigation time for the common stimulus.

#### Calcium signal data analysis

Calcium signals were first fitted the 405 nm channel onto the 465 nm channel to de-trend signal bleaching and movement artifacts, according to the manufacturer’s protocol (https://github.com/tjd2002/tjd-shared-code/blob/master/matlab/photometry/FP_normalize.m). Next, the de-trended signal was aligned to the video recording using the timestamps recorded by the digital port of the RZ10× system.

For calculating the mean signal amplitude over time, we first calculated the area under the curve (AUC) for each session in time bins of five seconds (using the original de-trended signal which is samples at 1017.3Hz) and then averaged the signals across all sessions. We then z-scored the AUC values for the 5-minutes of encounter using the average and standard deviation of the one-minute proceeding the encounter starting point (minute 4 of the pre-encounter).

#### Synchronization of physiological data to behavioral events

We used a custom-made code, termed DeepPhenotyping, to visualize and synchronize the de-trended signals and behavioral events, as previously described [90, 91]. The software considered the relevant sample rates of the video recording and the calcium signal recording, aligned the signal to the beginning of each behavioral event and normalized it using z*-*score (0.5 s bins), with the pre-event period of -4 s to -1 s as baseline. For calculating the subjects’ investigation probability of a given stimulus between different optic stimulation conditions, the investigation time for the relevant stimulus was synchronized to the onset of all bouts (regardless of type) which is also the onset of the optical stimulation in conditions where optic stimulation was delivered. Derived investigation probabilities were then normalized to the maximum (divided by P_max_ of 1) and used directly for statistical Analysis.

#### Synchronization of stimulus investigation to calcium transients

For calculating the probability of social investigation following significant negative or positive changes in calcium activity, we first segmented the de-trended calcium signal into 0.2-s time bins. For each bin, the signal was z-scored within an 8-s window (−4 s to +4 s relative to the bin onset) with the preceding 4 s (−4 s to 0 s) serving as baseline. The mean z-score during the 4 s following each bin was then compared to baseline activity. Time bins showing a significant deviation (Z ≥ +1.96 or Z ≤ -1.96) were classified as either positive or negative calcium events. Following the detection of a positive or negative event, a 6-s interval was imposed before the next sampling bin to ensure that detected events were distinct and did not reflect overlapping changes in the recorded signal at adjacent time points. All identified positive and negative events were subsequently aligned with the animal’s investigation time of a given stimulus, and the probability of investigation was calculated in 8-s time windows centered on the event onset. The mean investigation probability during the −4 to 0 s interval served as the baseline for z-scoring post-event (0 to 4 s) investigation probabilities. Z-scored investigation probabilities for positive and negative calcium transients were then averaged across all events pooled within each session. The AUC in the 0 to 4 s time window was computed to quantify differences in investigation probability following positive versus negative calcium transients, as well as to assess deviations from baseline (0) within each event type.

### Statistical Analysis

All statistical results and parameters are presented in Table S1, organized according to the corresponding figures. Sample sizes for all behavioral experiments were determined based on previously published power calculations [89]. Statistical analyses were performed using SPSS v27.0 (IBM) and GraphPad Prism v10.1.2 (GraphPad Software). The Kolmogorov–Smirnov and Shapiro–Wilk tests were used to assess the normality of the dependent variables. For normally distributed data, one-sample *t*-tests were used to compare group means to a fixed reference value (0 in this case). Two-tailed paired *t*-tests were used to compare conditions or stimuli within the same group, and two-tailed independent *t*-tests were used to compare variables between distinct groups. When normality assumptions were violated, nonparametric alternatives were applied. The Wilcoxon signed-rank test was used to compare group medians to a set value (0) or between conditions within the same group, and the Mann–Whitney *U* test was used to compare conditions between groups. For comparisons involving multiple factors, analyses of variance (ANOVA) were conducted according to the experimental design. Mixed-model (MM) ANOVA was applied when data included both within-subject (repeated-measures) and between-subject factors. This model included one random effect (subject ID), one within-subject effect, one between-subject effect, and their interaction. Two-way between-subjects ANOVA was used for comparisons between groups involving two independent factors, whereas two-way repeated-measures (RM) ANOVA was used for comparisons within groups across multiple conditions or variables. The latter model included one random effect (subject ID), two within-subject effects, and their interaction. When the assumption of sphericity required for ANOVA was violated, the Greenhouse–Geisser correction was applied. Whenever significant main effects or interactions were detected, post hoc Student’s *t*-tests with Holm–Šidák correction were conducted. For repeated-measures data violating normality assumptions, Friedman’s nonparametric test was applied, followed by post hoc Dunn’s paired comparisons. For between-group comparisons involving more than two levels, the Kruskal–Wallis test was used, followed by Dunn’s multiple-comparisons post hoc tests. Statistical significance was set at *p* < 0.05.

## References

1. Chen, P. and W. Hong, Neural Circuit Mechanisms of Social Behavior. Neuron, 2018. 98(1): p. 16–30.

2. Gangopadhyay, P., et al., Prefrontal–amygdala circuits in social decision-making. Nature neuroscience, 2021. 24(1): p. 5–18.

3. Kietzman, H.W. and S.L. Gourley, How social information impacts action in rodents and humans: the role of the prefrontal cortex and its connections. Neuroscience & Biobehavioral Reviews, 2023. 147: p. 105075.

4. Cao, W., H. Li, and J. Luo, Prefrontal cortical circuits in social behaviors: an overview. Journal of Zhejiang University-SCIENCE B, 2024. 25(11): p. 941–955.

5. Bicks, L.K., et al., Prefrontal Cortex and Social Cognition in Mouse and Man. Front Psychol, 2015. 6: p. 1805.

6. Ko, J., Neuroanatomical Substrates of Rodent Social Behavior: The Medial Prefrontal Cortex and Its Projection Patterns. Front Neural Circuits, 2017. 11: p. 41.

7. Lee, E., et al., Enhanced neuronal activity in the medial prefrontal cortex during social approach behavior. Journal of Neuroscience, 2016. 36(26): p. 6926–6936.

8. Yashima, J., T. Uekita, and T. Sakamoto, The prelimbic cortex but not the anterior cingulate cortex plays an important role in social recognition and social investigation in mice. PloS one, 2023. 18(4): p. e0284666.

9. Kuga, N., et al., Prefrontal-amygdalar oscillations related to social behavior in mice. eLife, 2022. 11: p. e78428.

10. Li, H., et al., Brain circuits that regulate social behavior. Molecular Psychiatry, 2025: p. 1–17.

11. van den Bos, W. and B. Güroğlu, The role of the ventral medial prefrontal cortex in social decision making. Journal of Neuroscience, 2009. 29(24): p. 7631–7632.

12. Carlén, M., What constitutes the prefrontal cortex? Science, 2017. 358(6362): p. 478–482.

13. Anastasiades, P.G. and A.G. Carter, Circuit organization of the rodent medial prefrontal cortex. Trends in neurosciences, 2021. 44(7): p. 550–563.

14. Capuzzo, G. and S.B. Floresco, Prelimbic and infralimbic prefrontal regulation of active and inhibitory avoidance and reward-seeking. Journal of Neuroscience, 2020. 40(24): p. 4773–4787.

15. Giustino, T.F. and S. Maren, The role of the medial prefrontal cortex in the conditioning and extinction of fear. Frontiers in behavioral neuroscience, 2015. 9: p. 298.

16. Quirk, G.J. and D. Mueller, Neural mechanisms of extinction learning and retrieval. Neuropsychopharmacology, 2008. 33(1): p. 56–72.

17. Kietzman, H.W., et al., Social incentivization of instrumental choice in mice requires amygdala-prelimbic cortex-nucleus accumbens connectivity. Nature Communications, 2022. 13(1): p. 4768.

18. Glanzberg, J.T., et al., Individual differences in prelimbic neural representation of food and cocaine seeking. Cell reports, 2024. 43(12).

19. Frost, N.A., et al., Context-invariant socioemotional encoding by prefrontal ensembles. Nature Communications, 2025. 16(1): p. 5455.

20. Isaac, J., et al., Sex differences in neural representations of social and nonsocial reward in the medial prefrontal cortex. Nature Communications, 2024. 15(1): p. 8018.

21. Klune, C.B., B. Jin, and L.A. DeNardo, Linking mPFC circuit maturation to the developmental regulation of emotional memory and cognitive flexibility. Elife, 2021. 10: p. e64567.

22. Ebstein, R.P., et al., Genetics of human social behavior. Neuron, 2010. 65(6): p. 831–844.

23. McGraw, L.A. and L.J. Young, The prairie vole: an emerging model organism for understanding the social brain. Trends in neurosciences, 2010. 33(2): p. 103–109.

24. Netser, S., et al., Distinct dynamics of social motivation drive differential social behavior in laboratory rat and mouse strains. Nat Commun, 2020. 11(1): p. 5908.

25. Jabarin, R., et al., Distinct prelimbic cortex neuronal responses to emotional states of others drive emotion recognition in adult mice. Current Biology, 2025. 35(5): p. 994–1011. e8.

26. Jabarin, R., et al., Modulation of social investigation by anterior hypothalamic nucleus rhythmic neural activity. Iscience, 2023. 26(2).

27. Hyman, J.M., et al., Contextual encoding by ensembles of medial prefrontal cortex neurons. Proceedings of the National Academy of Sciences, 2012. 109(13): p. 5086–5091.

28. Walton, M.E., et al., Separable learning systems in the macaque brain and the role of orbitofrontal cortex in contingent learning. Neuron, 2010. 65(6): p. 927–939.

29. Marton, T.F., et al., Roles of prefrontal cortex and mediodorsal thalamus in task engagement and behavioral flexibility. Journal of Neuroscience, 2018. 38(10): p. 2569–2578.

30. Ragozzino, M.E., The contribution of the medial prefrontal cortex, orbitofrontal cortex, and dorsomedial striatum to behavioral flexibility. Annals of the New York academy of sciences, 2007. 1121(1): p. 355–375.

31. Gabriel, C.J., et al., Transformations in prefrontal ensemble activity underlying rapid threat avoidance learning. Current Biology, 2025. 35(5): p. 1128–1136. e4.

32. Schoenbaum, G., A.A. Chiba, and M. Gallagher, Neural encoding in orbitofrontal cortex and basolateral amygdala during olfactory discrimination learning. Journal of Neuroscience, 1999. 19(5): p. 1876–1884.

33. Rudebeck, P.H., et al., Separate neural pathways process different decision costs. Nature neuroscience, 2006. 9(9): p. 1161–1168.

34. Sul, J.H., et al., Distinct roles of rodent orbitofrontal and medial prefrontal cortex in decision making. Neuron, 2010. 66(3): p. 449–460.

35. Choi, K., et al., Distributed processing for value-based choice by prelimbic circuits targeting anterior-posterior dorsal striatal subregions in male mice. Nature Communications, 2023. 14(1): p. 1920.

36. Niedringhaus, M. and E.A. West, Prelimbic cortex neural encoding dynamically tracks expected outcome value. Physiology & behavior, 2022. 256: p. 113938.

37. Kentrop, J., et al., Pro-social preference in an automated operant two-choice reward task under different housing conditions: Exploratory studies on pro-social decision making. Developmental Cognitive Neuroscience, 2020. 45: p. 100827.

38. Martin, L. and E. Iceberg, Quantifying social motivation in mice using operant conditioning. Journal of visualized experiments: JoVE, 2015(102): p. 53009.

39. Cheng, Y., et al., Asymmetric Social Representations in the Prefrontal Cortex for Cooperative Behavior. bioRxiv, 2025.

40. Alabi, O.O., M.P. Fortunato, and M.V. Fuccillo, Behavioral paradigms to probe individual mouse differences in value-based decision making. Frontiers in neuroscience, 2019. 13: p. 50.

41. Haber, S.N. and B. Knutson, The reward circuit: linking primate anatomy and human imaging. Neuropsychopharmacology, 2010. 35(1): p. 4–26.

42. Ali, F. and A.C. Kwan, Interpreting in vivo calcium signals from neuronal cell bodies, axons, and dendrites: a review. Neurophotonics, 2020. 7(1): p. 011402–011402.

43. Simpson, E.H., et al., Lights, fiber, action! A primer on in vivo fiber photometry. Neuron, 2024. 112(5): p. 718–739.

44. Legaria, A.A., J.A. Licholai, and A.V. Kravitz, Fiber photometry does not reflect spiking activity in the striatum. BioRxiv, 2021: p. 2021.01. 20.427525.

45. Klavir, O., et al., Manipulating fear associations via optogenetic modulation of amygdala inputs to prefrontal cortex. Nature neuroscience, 2017. 20(6): p. 836–844.

46. Levy, D.R., et al., Dynamics of social representation in the mouse prefrontal cortex. Nat Neurosci, 2019. 22(12): p. 2013–2022.

47. Yu, Y.H., et al., Optogenetic stimulation in the medial prefrontal cortex modulates stimulus valence from rewarding and aversive to neutral states. Frontiers in Psychiatry, 2023. 14: p. 1119803.

48. Yizhar, O., Optogenetic insights into social behavior function. Biological psychiatry, 2012. 71(12): p. 1075–1080.

49. Yizhar, O., et al., Optogenetics in neural systems. Neuron, 2011. 71(1): p. 9–34.

50. Tan, T., et al., Neural circuits and activity dynamics underlying sex-specific effects of chronic social isolation stress. Cell Rep, 2021. 34(12): p. 108874.

51. Bromberg-Martin, E.S., M. Matsumoto, and O. Hikosaka, Dopamine in motivational control: rewarding, aversive, and alerting. Neuron, 2010. 68(5): p. 815–834.

52. Coenen, V.A., et al., The anatomy of the human medial forebrain bundle: ventral tegmental area connections to reward-associated subcortical and frontal lobe regions. NeuroImage: Clinical, 2018. 18: p. 770–783.

53. Felix-Ortiz, A.C., et al., Bidirectional modulation of anxiety-related and social behaviors by amygdala projections to the medial prefrontal cortex. Neuroscience, 2016. 321: p. 197–209.

54. Scheggia, D. and F. Papaleo, Social Neuroscience: Rats Can Be Considerate to Others. Curr Biol, 2020. 30(6): p. R274–R276.

55. Yang, S.-S., et al., Prefrontal GABAergic interneurons gate long-range afferents to regulate prefrontal cortex-associated complex behaviors. Frontiers in neural circuits, 2021. 15: p. 716408.

56. McGarry, L.M. and A.G. Carter, Inhibitory gating of basolateral amygdala inputs to the prefrontal cortex. Journal of Neuroscience, 2016. 36(36): p. 9391–9406.

57. Gu, X. and J.P. Johansen, Prefrontal encoding of an internal model for emotional inference. Nature, 2025: p. 1–13.

58. Ruff, C.C. and E. Fehr, The neurobiology of rewards and values in social decision making. Nature Reviews Neuroscience, 2014. 15(8): p. 549–562.

59. Chib, V.S., et al., Evidence for a common representation of decision values for dissimilar goods in human ventromedial prefrontal cortex. Journal of Neuroscience, 2009. 29(39): p. 12315–12320.

60. Levy, D.J. and P.W. Glimcher, The root of all value: a neural common currency for choice. Current opinion in neurobiology, 2012. 22(6): p. 1027–1038.

61. Fujimoto, A., E.A. Murray, and P.H. Rudebeck, Interaction between decision-making and interoceptive representations of bodily arousal in frontal cortex. Proceedings of the National Academy of Sciences, 2021. 118(35): p. e2014781118.

62. Lui, J.H., et al., Differential encoding in prefrontal cortex projection neuron classes across cognitive tasks. Cell, 2021. 184(2): p. 489–506 e26.

63. Choi, T.-Y., et al., Distinct prefrontal projection activity and transcriptional state conversely orchestrate social competition and hierarchy. Neuron, 2024. 112(4): p. 611–627. e8.

64. Liu, Y., et al., A molecularly defined mPFC-BLA circuit specifically regulates social novelty preference. Science Advances, 2025. 11(17): p. eadt9008.

65. Ito, R., T.W. Robbins, and B.J. Everitt, Differential control over cocaine-seeking behavior by nucleus accumbens core and shell. Nature neuroscience, 2004. 7(4): p. 389–397.

66. Lai, C.-H., et al., Decoding the hidden variabilities in mPFC descending pathways across emotional states. eLife, 2025. 14.

67. Murugan, M., et al., Combined social and spatial coding in a descending projection from the prefrontal cortex. Cell, 2017. 171(7): p. 1663–1677. e16.

68. Kietzman, H.W., et al., Neuronal ensembles in the amygdala allow social information to motivate later decisions. Journal of Neuroscience, 2024. 44(16).

69. Janak, P.H. and K.M. Tye, From circuits to behaviour in the amygdala. Nature, 2015. 517(7534): p. 284–292.

70. Ferguson, B.R. and W.-J. Gao, PV interneurons: critical regulators of E/I balance for prefrontal cortex-dependent behavior and psychiatric disorders. Frontiers in neural circuits, 2018. 12: p. 37.

71. Scheggia, D., et al., Somatostatin interneurons in the prefrontal cortex control affective state discrimination in mice. Nature neuroscience, 2020. 23(1): p. 47–60.

72. Lam, N.H., et al., Effects of altered excitation-inhibition balance on decision making in a cortical circuit model. Journal of Neuroscience, 2022. 42(6): p. 1035–1053.

73. Yizhar, O., et al., Neocortical excitation/inhibition balance in information processing and social dysfunction. Nature, 2011. 477(7363): p. 171–178.

74. Liu, L., et al., Cell type–differential modulation of prefrontal cortical GABAergic interneurons on low gamma rhythm and social interaction. Science advances, 2020. 6(30): p. eaay4073.

75. Rubenstein, J.L. and M.M. Merzenich, Model of autism: increased ratio of excitation/inhibition in key neural systems. Genes, Brain and Behavior, 2003. 2(5): p. 255–267.

76. Selimbeyoglu, A., et al., Modulation of prefrontal cortex excitation/inhibition balance rescues social behavior in CNTNAP2-deficient mice. Science translational medicine, 2017. 9(401): p. eaah6733.

77. Sierra-Mercado, D., N. Padilla-Coreano, and G.J. Quirk, Dissociable roles of prelimbic and infralimbic cortices, ventral hippocampus, and basolateral amygdala in the expression and extinction of conditioned fear. Neuropsychopharmacology, 2011. 36(2): p. 529–538.

78. Vidal-Gonzalez, I., et al., Microstimulation reveals opposing influences of prelimbic and infralimbic cortex on the expression of conditioned fear. Learning & memory, 2006. 13(6): p. 728–733.

79. Quirk, G.J., et al., Stimulation of medial prefrontal cortex decreases the responsiveness of central amygdala output neurons. Journal of Neuroscience, 2003. 23(25): p. 8800–8807.

80. Riaz, S., et al., Prelimbic and infralimbic cortical inactivations attenuate contextually driven discriminative responding for reward. Scientific reports, 2019. 9(1): p. 3982.

81. Gutman, A.L., et al., The infralimbic and prelimbic cortices contribute to the inhibitory control of cocaine -seeking behavior during a discriminative stimulus task in rats. Addiction biology, 2017. 22(6): p. 1719–1730.

82. Kwan, A.C., What can population calcium imaging tell us about neural circuits? Journal of neurophysiology, 2008. 100(6): p. 2977–2980.

83. Netser, S., et al., A novel system for tracking social preference dynamics in mice reveals sex-and strain-specific characteristics. Molecular autism, 2017. 8(1): p. 53.

84. Aravanis, A.M., et al., An optical neural interface: in vivo control of rodent motor cortex with integrated fiberoptic and optogenetic technology. Journal of neural engineering, 2007. 4(3): p. S143.

85. Cela, E., et al., An optogenetic kindling model of neocortical epilepsy. Scientific Reports, 2019. 9(1): p. 5236.

86. Britt, J.P., R.A. McDevitt, and A. Bonci, Use of channelrhodopsin for activation of CNS neurons. Current protocols in neuroscience, 2012. 58(1): p. 2.16.1–2.16.19.

87. Lin, J.Y., A user’s guide to channelrhodopsin variants: features, limitations and future developments. Experimental physiology, 2011. 96(1): p. 19–25.

88. Netser, S., et al., A novel system for tracking social preference dynamics in mice reveals sex- and strain-specific characteristics. Mol Autism, 2017. 8: p. 53.

89. Netser, S., et al., A system for tracking the dynamics of social preference behavior in small rodents. Journal of Visualized Experiments (JoVE), 2019(153): p. e60336.

90. Barbier, M., et al., Altered Neural Activity in the Mesoaccumbens Pathway Underlies Impaired Social Reward Processing in Shank3 -Deficient Rats. Advanced Science, 2025. 12(17): p. 2414813.

91. Mohapatra, A.N., et al., Impaired emotion recognition in Cntnap2-deficient mice is associated with hyper-synchronous prefrontal cortex neuronal activity. Molecular Psychiatry, 2025. 30(4): p. 1440–1452.

