## Supplementary figures for "Inhibition governs preference encoding in medial prefrontal cortex pyramidal neurons during a binary social choice in mice"

**Document Type:** Supplementary Figures and Legends

This file contains all supplementary figures and their corresponding legends (Figures S1–S9), referenced in the main manuscript titled "Inhibition governs stimulus preference encoding in mPFC pyramidal neurons during social choice." All figure numbers correspond to their citations within the main text.

### Table of Contents

| Figure | Title / Description |
|---|---|
| *Figure S1.* | Behavioral parameters of exploration, locomotion, and preference across four social discrimination tasks |
| *Figure S2.* | Calcium signal dynamics across social contexts |
| *Figure S3.* | Baseline investigation and mPFC activity during pre-encounter stages |
| *Figure S4.* | Non-transitional bout activity across discrimination tasks |
| *Figure S5.* | Lack of subregional specificity in mPFC activity during social choice |
| *Figure S6.* | Workflow of calcium event–behavior probability analysis |
| *Figure S7.* | Positive calcium events and investigation behavior |
| *Figure S8.* | Behavioral metrics across optogenetic stimulation conditions |
| *Figure S9.* | mPFC Pyramidal Neurons show no responses during non-transitional bouts in the social fear conditioning paradigm |
| *Figure S10.* | Activity of mPFC-to-NAc and mPFC-to-BLA neurons during non-transitional bouts. |

All data supporting these figures are available in the supplementary tables accompanying the manuscript.

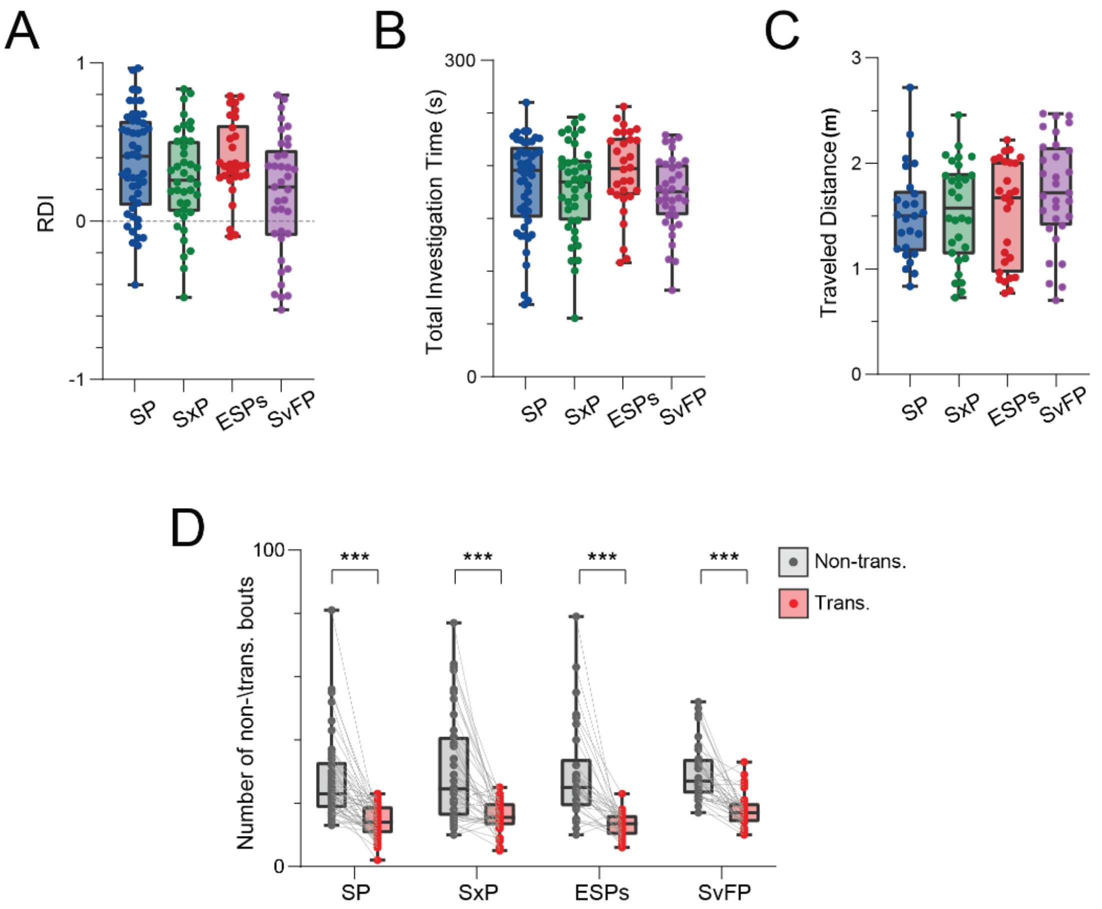


**Figure S1. Behavioral parameters of exploration, locomotion, and preference of subject mice across the four social discrimination tasks**

- **A.** Median (in box plot*) relative discrimination index (RDI) for the preferred stimulus during the 5 min encounter period of the SP, SxP, ESPs, and SvFP tasks.
- **B.** Median (in box plot*) total investigation time during the 5-minute encounter period in each of the four tasks.
- **C.** Median (in box plot*) traveled distance during the 5-minute encounter period in each of the four tasks.
- **D.** Median (in box plot*) sum of non-transitional (in gray) and transitional (in red) bouts during the 5 min encounter period in each of the four tasks.

***$p<0.001$, Wilcoxon signed-rank test with Holm-Šídák correction for multiple comparisons following detection of main effects by ANOVA.

Box plot represents 25 to 75 percentiles of the distribution, while the bold line is the median of the distribution. Whiskers represent the smallest and largest values in the distribution.

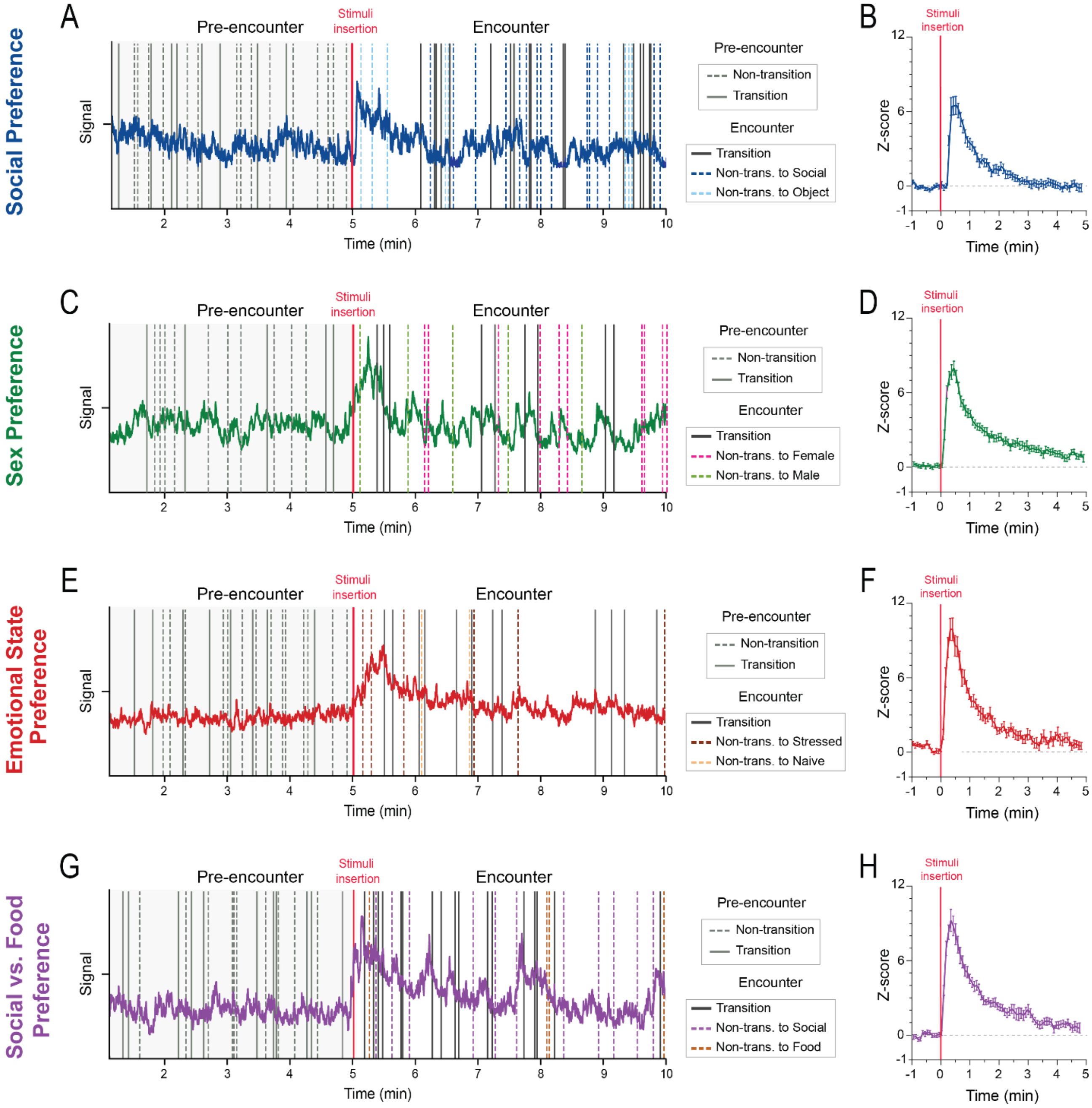

Social Preference
A
Pre-encounter
Stimuli insertion
Encounter
Signal
Time (min)
Pre-encounter
Non-transition
Transition
Encounter
Transition
Non-trans. to Social
Non-trans. to Object
B
Z-score
Time (min)
Sex Preference
C
Non-trans. to Female
Non-trans. to Male
D
Emotional State Preference
E
Non-trans. to Stressed
Non-trans. to Naive
F
Social vs. Food Preference
G
Non-trans. to Social
Non-trans. to Food
H

**Figure S2. Social encounters of varying contexts elicit a similar pattern of calcium signal activity in pyramidal neurons of the mPFC**

**A.** An example trace of normalized calcium signals recorded across the pre-encounter (before stimulus insertion) and encounter (after stimulus insertion) stages of an SP session. Onsets of transitional bouts are denoted by filled lines (colored in gray in the pre-encounter stage, and in black in the encounter stage). Onsets of non-transitional bouts are denoted in dashed lines (colored in gray in the pre-encounter stage, and colored according to stimulus in the encounter stage).

**B.** Mean (±SEM, 100-ms bins) z-scored amplitude of the calcium signal over time, averaged across all SP sessions, with the last minute of the pre-encounter stage serving as baseline. The red line indicates the time of stimuli insertion and the start of the encounter stage.

**C-D.** As in **A-B**, for the SxP task.

**E-F.** As in **A-B**, for the ESPs task.

**G-H.** As in **A-B**, for the SvFP task.

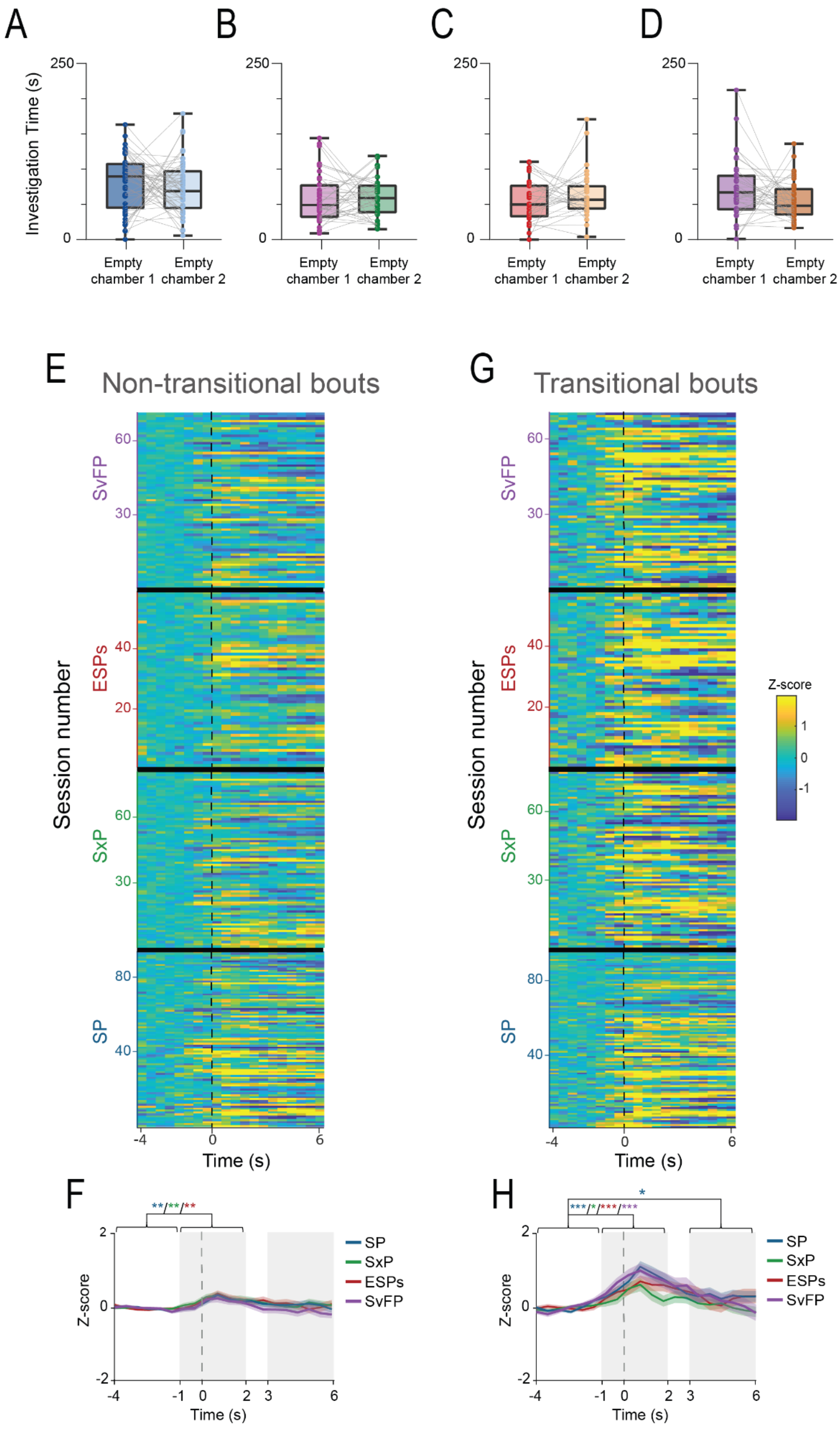
A
B
C
D
Investigation Time (s)
250
0
Empty chamber 1
Empty chamber 2
E
Non-transitional bouts
G
Transitional bouts
Session number
SvFP
ESPs
SxP
SP
Z-score
1
0
-1
Time (s)
-4
0
6
F
H
SP
SxP
ESPs
SvFP
Z-score
2
-2
-1
2
3

**Figure S3. Baseline investigation and neural activity during the pre-encounter stage**

**A.** Median time dedicated by subject animals during the 5 minutes of the pre-encounter stage to investigate the empty chambers later used to house the social/preferred stimulus (chamber 1, dark blue) and the object/non-preferred (chamber 2, light blue) in the encounter stage of all SP sessions. Each grey line connects data points of the same session.

**B-D.** As in **A**, for the SxP, ESPs, and SvFP tasks, respectively.

**E.** Heat-map (see color code to the right of **G**) of the z-scored calcium signals at the beginning (first six seconds) of bout, averaged across all non-transitional bouts towards the empty chambers during the 5 min pre-encounter stage of the SP, SxP, ESPs, and SvFP task, from the bottom up (each row corresponds to a single session), using 0.5-s bins. Time ‘0’ represents the beginning of the bout.

**F.** Superimposed traces of the mean (±SEM) z-scored calcium signals shown in **E,** averaged across all SP (in dark blue), SxP (in green), ESPs (in red), and SvFP (in purple) sessions. Gray-shaded areas represent two distinct time windows of the event, which are used for statistical analysis. A significant effect was found for time ($p<0.05$ in SxP and SvFP) by one-way RM ANOVA.

**G-H.** As in **E-F**, for transitional bouts. A significant effect was found for time ($p<0.001$ in SP and SvF; $p<0.01$ in SxP and ESPs) by one-way RM ANOVA.

*$p<0.05$, **$p$<0.01, ***$p$<0.001, *post-hoc* paired t-test with Holm-Šídák correction for multiple comparisons following the detection of main effects by ANOVA.

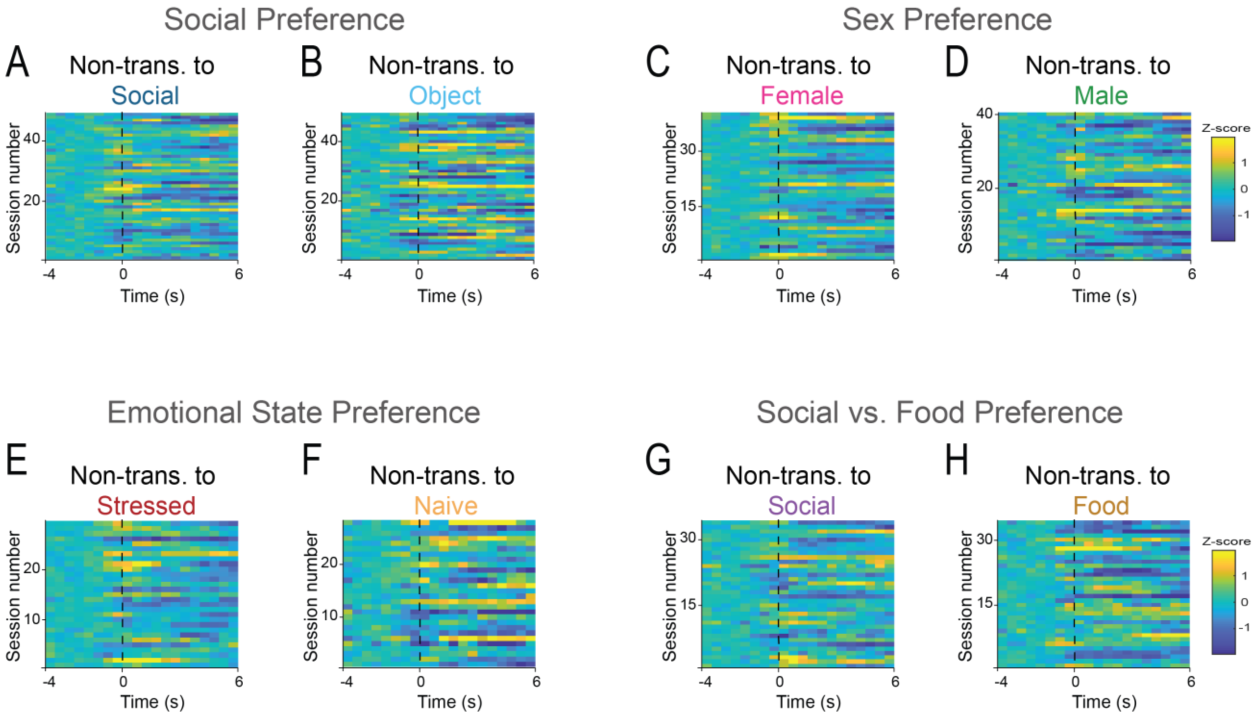


**Figure S4. Non-transitional bout activity across the four social discrimination tasks**

- **A.** Heat-map (see color code to the right of **D**) of the z-scored calcium signals at the beginning (first six seconds) of bout, averaged across all non-transitional bouts towards the social stimulus during the 5-minute encounter stage of the SP sessions, using 0.5-s bins. Time '0' represents the beginning of the bout.
- **B.** As in **A**, for non-transitional bouts to the object.
- **C-D**. As in **A-B**, for non-transitional bouts to the female and the male stimuli, respectively, in the SxP task.
- **E-F**. As in **A-B**, for non-transitional bouts to the stressed and the naive stimuli, respectively, in the ESPs task.
- **G-H**. As in **A-B**, for non-transitional bouts to the social stimulus and the food, respectively, in the SvFP task.

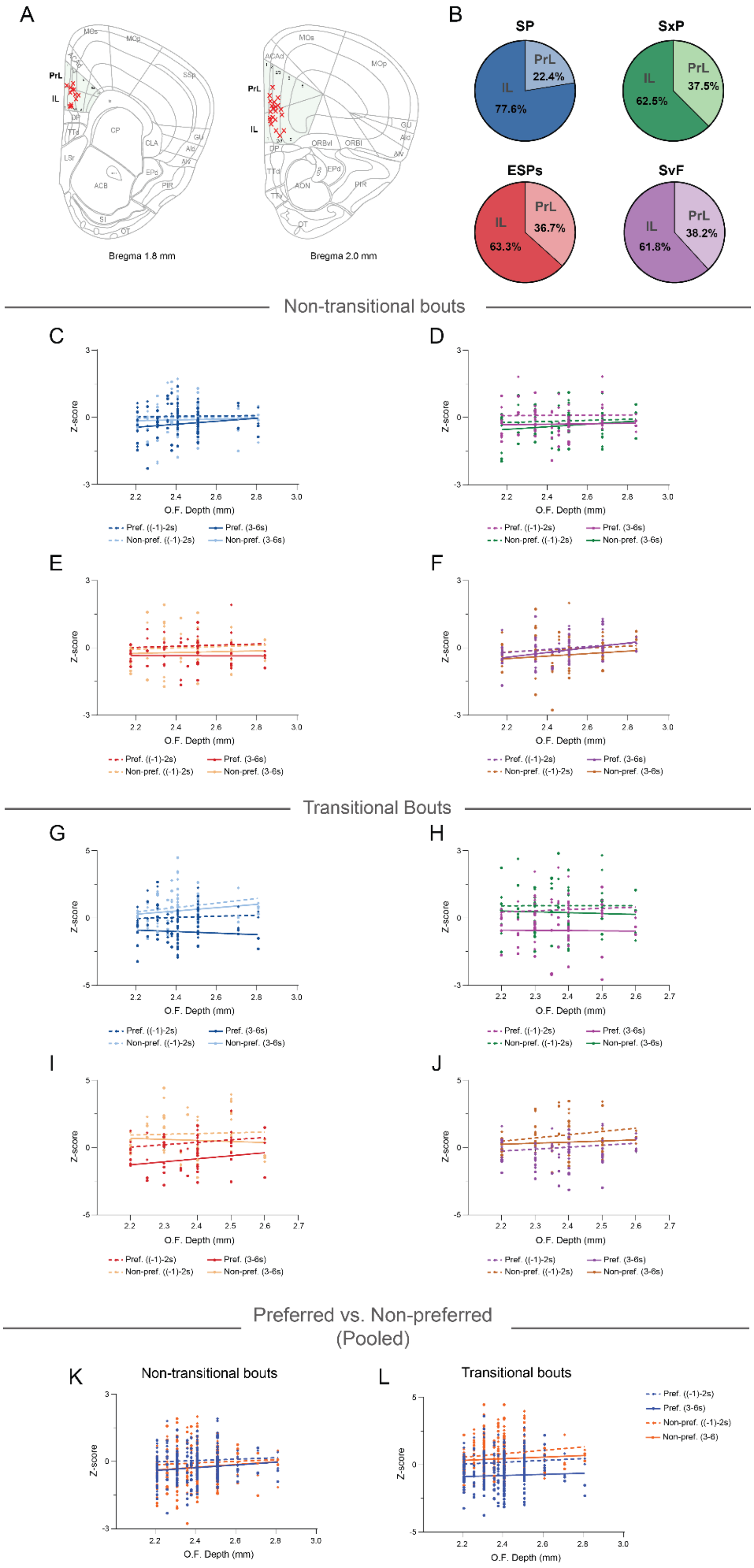

A
PrL
IL
Bregma 1.8 mm
Bregma 2.0 mm
B
SP
PrL
22.4%
IL
77.6%
SxP
PrL
37.5%
IL
62.5%
ESPs
PrL
36.7%
IL
63.3%
SvF
PrL
38.2%
IL
61.8%
Non-transitional bouts
C
D
E
F
Z-score
O.F. Depth (mm)
Pref. ((-1)-2s)
Pref. (3-6s)
Non-pref. ((-1)-2s)
Non-pref. (3-6s)
Transitional Bouts
G
H
I
J
Preferred vs. Non-preferred
(Pooled)
K
Non-transitional bouts
L
Transitional bouts
Non-pref. (3-6)

**Figure S5. Lack of subregional specificity in mPFC activity during social choice**

**A.** A schematic representation of a coronal forebrain slice at 2mm (right slice) and 1.8mm (left slice) anterior to Bregma, showing the exact location of the optic fiber tip within the mPFC of each subject animal across all fiber photometry experiments (n=35 animals), marked in red X.

**B.** Pie charts depicting the percentage of recorded sessions from animals with optic fiber tip in the PrL and IL regions in the SP, SxP, ESPs, and SvFP tasks.

**C.** Correlation between optic fiber tip depth (the dorsoventral location) and the averaged z-score for the recorded calcium signal for non-transitional bouts to the preferred (social) and the non-preferred (object) stimuli across all SP sessions in two different time windows relative to the onset of the non-transitional bout (-1to2s, and 3to6s).

**D-F.** As in **C**, for SxP, ESPs, and SvFP sessions, respectively.

**G-J.** As in **C-F**, for transitional bouts in the SP, SxP, ESPs, and SvFP sessions, respectively.

**K.** As in **C**, for non-transitional bouts to the preferred and the non-preferred stimuli, pooled from all four tasks.

**L.** As in **K**, for transitional bouts.

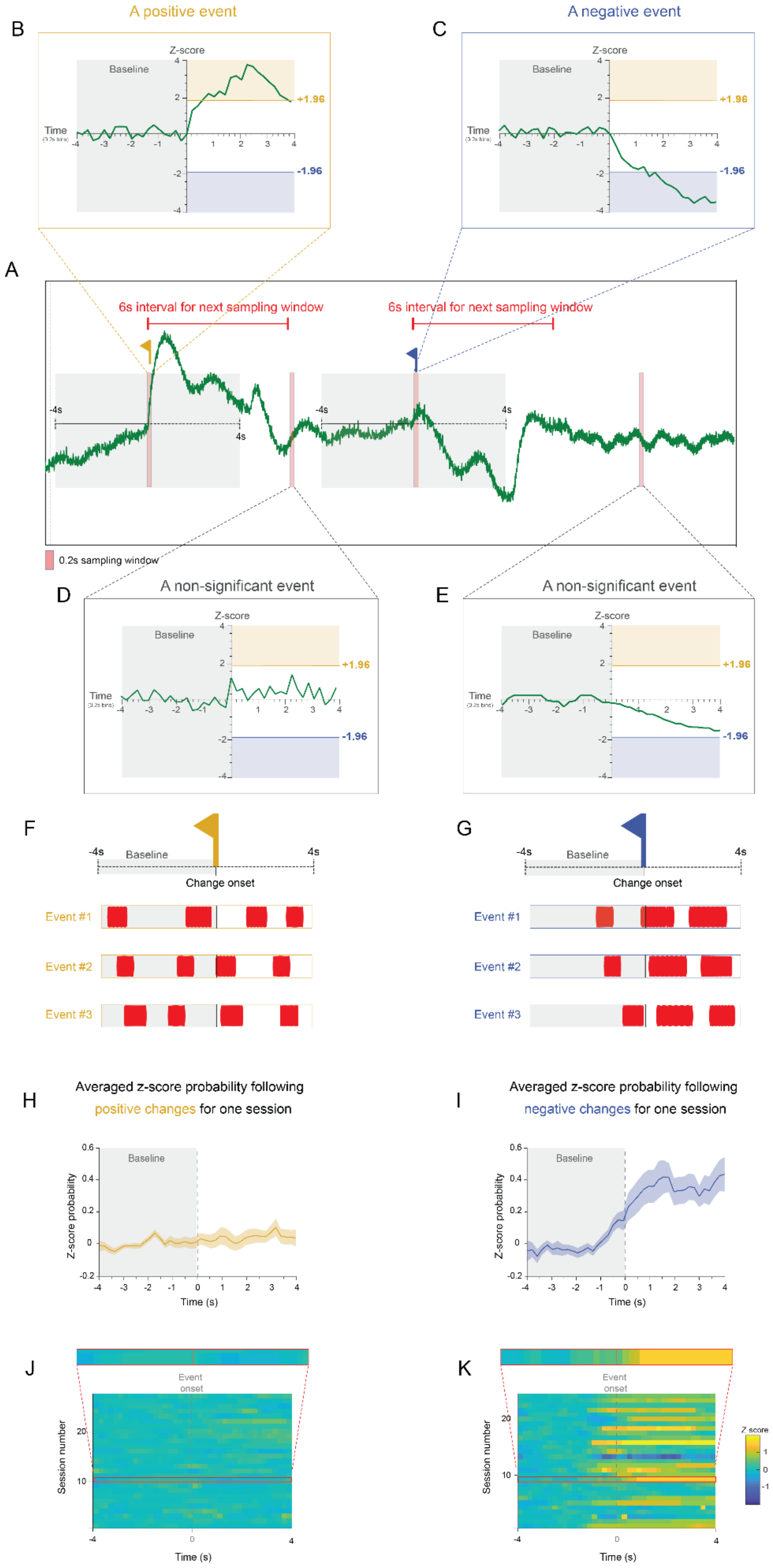

A positive event
A negative event
B
C
A
6s interval for next sampling window
6s interval for next sampling window
0.2s sampling window
D
E
A non-significant event
A non-significant event
Z-score
Baseline
Time
+1.96
-1.96
F
G
Baseline
Change onset
Event #1
Event #2
Event #3
H
I
Averaged z-score probability following positive changes for one session
Averaged z-score probability following negative changes for one session
Z-score probability
Time (s)
J
K
Event onset
Session number
Time (s)
Z score

**Figure S6. Schematic and workflow of the calcium event–behavior probability analysis**

- **A.** A 30s segment of a normalized calcium signal trace. The examined trace is first segmented into 0.2-s bins (represented by the pink bar, “A sampling window”). For each bin, an 8-s analysis window was established, consisting of a 4-s baseline period immediately preceding the bin’s onset (–4 s to 0 s) and a 4-s post-onset period (0 s to +4 s). Upon detection of a significant event (positive or negative), a 6-second interval was taken before the next sampling window.
- **B.** The mean z-scored signal during the 4-s post-onset window was compared to the 4-s baseline. The z-score average in the 4-s post-onset window was outside the range of +1.96 z-scores (the z-score threshold values for classifying significant events); therefore, the corresponding sampling window is flagged as a positive event in the trace in B represented by a yellow flag.
- **C.** As in **C**, for a negative change event in the calcium signal represented by a blue flag.
- **D-E.** As in **C**, for non-significant events with changes in z-score that do not cross the $\pm$1.96 z-score thresholds.
- **F.** The subject’s investigation of the stimulus (red bars) was aligned for each identified positive event in an 8-second window (4 seconds before and after the onset of the change event). The probability of stimulus investigation is then calculated for each event type in 0.2-second bins.
- **G.** As in **F**, for negative events.
- **H.** Superimposed trace of the stimulus investigation probability 4 seconds following positive events, z-scored to the probability in the -4-to-0 window in 0.2s bins, and averaged across all positive events in the session. Time ‘0’ indicates the onset of the positive change event, with the shaded area representing the standard error of the mean ($\pm$SEM).
- **I.** As in **H**, for stimulus investigation probability following negative change events.
- **J.** Heat-map (see color code to the right) of the z-scored stimulus investigation probability following (4 seconds) positive change events in the calcium signal, averaged across all positive change events identified in the session, shown for all sessions of the task, with each row corresponding to a single session, with time ‘0’ representing the onset of positive change events. The row (session) corresponding to the trace in **I** is highlighted in red above the heatmap.
- **K.** As in **K**, for the stimulus investigation probability following negative change events in the calcium signal.

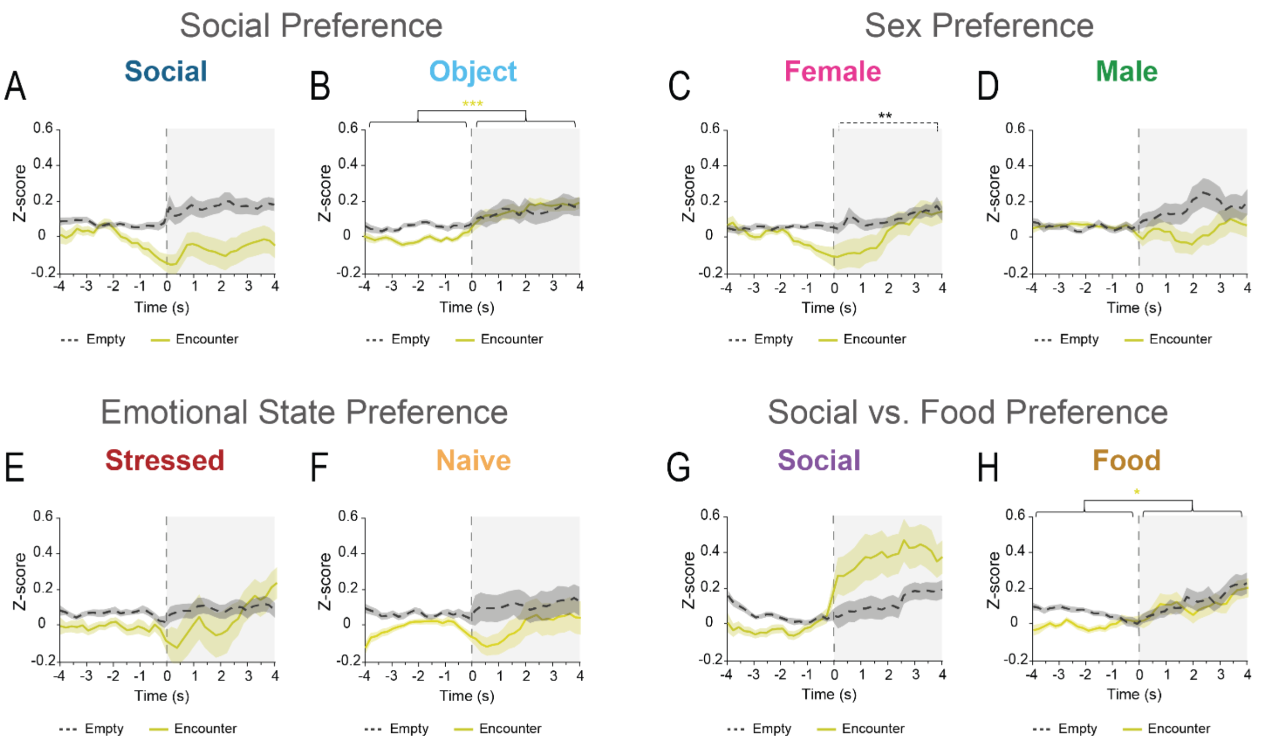


**Figure S7. Positive calcium events are not predictive of subsequent investigation behavior**

**A.** Superimposed traces of the mean (+SEM) social stimulus investigation probability for 4 seconds following the onset (time '0') of positive change events identified in the pre-encounter stage (black dashed line) and in the encounter stage (yellow filled line), z-scored to the probability in the -4to0 window in 0.2s bins, and averaged across all SP sessions.

**B.** As in **A-B**, for the object in SP sessions.

**C-D.** As in **A-B**, for the SxP task.

**E-F.** As in **A-B**, for the ESPs task.

**G-H.** As in **A-B**, for the SvFP task.

*$p<0.05$, **$p<0.01$, ***$p<0.001$, One-sample Wilcoxon signed rank test (within trace differences in time) or Wilcoxon signed rank test (differences between pre-encounter and encounter).

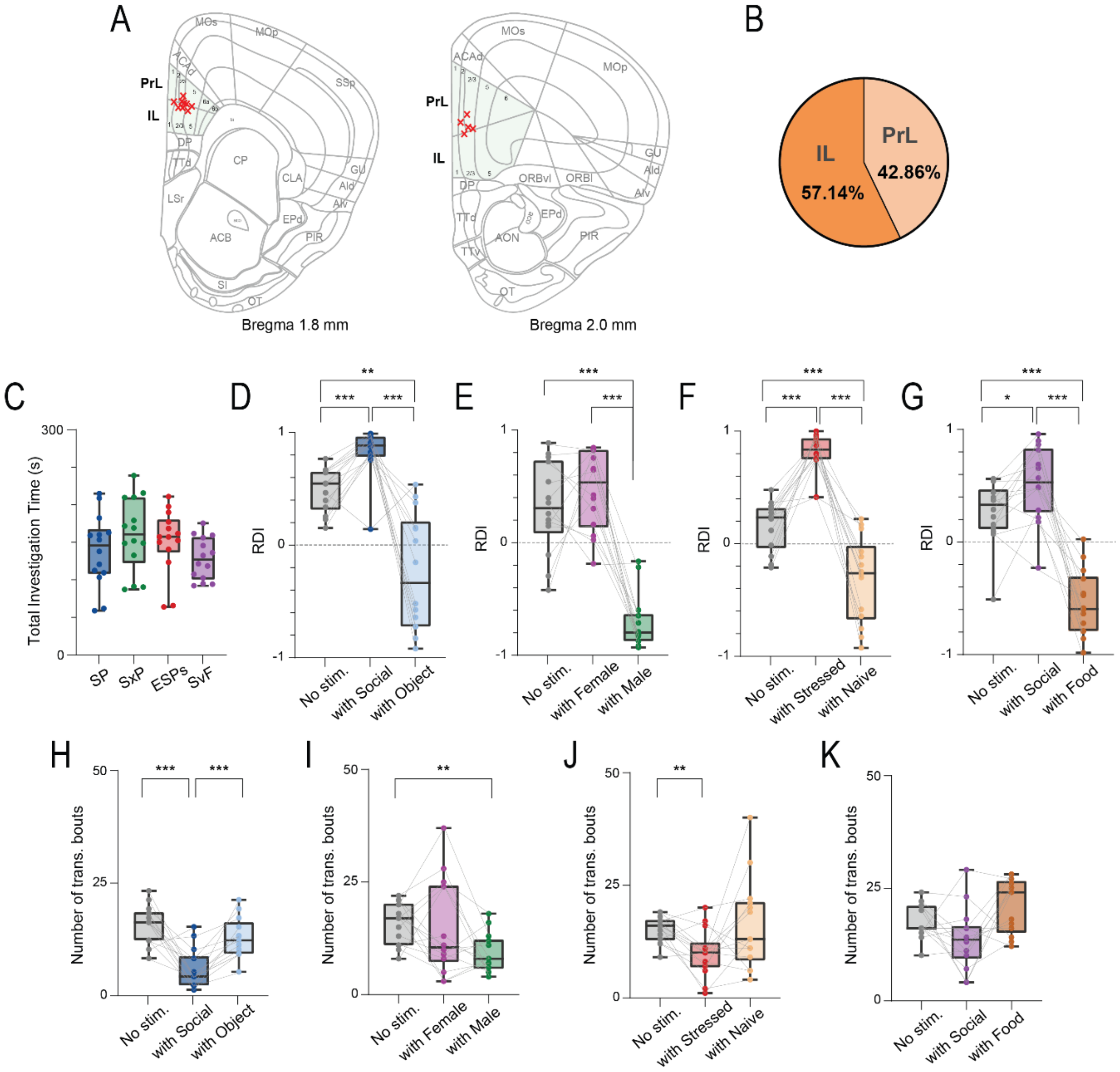

A
B
PrL
IL
Bregma 1.8 mm
Bregma 2.0 mm
PrL 42.86%
IL 57.14%
C
D
E
F
G
Total Investigation Time (s)
SP
SxP
ESPs
SvF
RDI
No stim.
with Social
with Object
with Female
with Male
with Stressed
with Naive
with Food
H
I
J
K
Number of trans. bouts

**Figure S8. Behavioral metrics across optogenetic stimulation conditions across the four social discrimination tasks**

**A.** A schematic representation of a coronal forebrain slice at 2mm (right slice) and 1.8mm (left slice) anterior to Bregma, showing the exact location of the optic fiber tip within the mPFC of each subject animal across all optogenetic experiments (n=14), marked in red X.

**B.** A pie chart depicting the percentage of animals with optic fiber tip in the PrL and IL sub-regions of the mPFC.

**C.** Median total investigation time for both stimuli in the no stimulation condition of each of the four tasks.

**D.** Median RDI for the typically preferred stimulus (social) in sessions from the three optic stimulation conditions of the SP task.

**E-G.** As in **B**, for the SxP, ESPs, and SvFP tasks, respectively.

**A.** Median sum of transitional bouts in sessions from the three optic stimulation conditions of the SP task.

**I-K.** As in **H**, for the SxP, ESPs, and SvFP tasks, respectively.

**p<0.05*, ***p*<0.01, ****p*<0.001, *post-hoc* paired t-test with Holm-Šídák correction for multiple comparisons following detection of main effects by RM ANOVA.

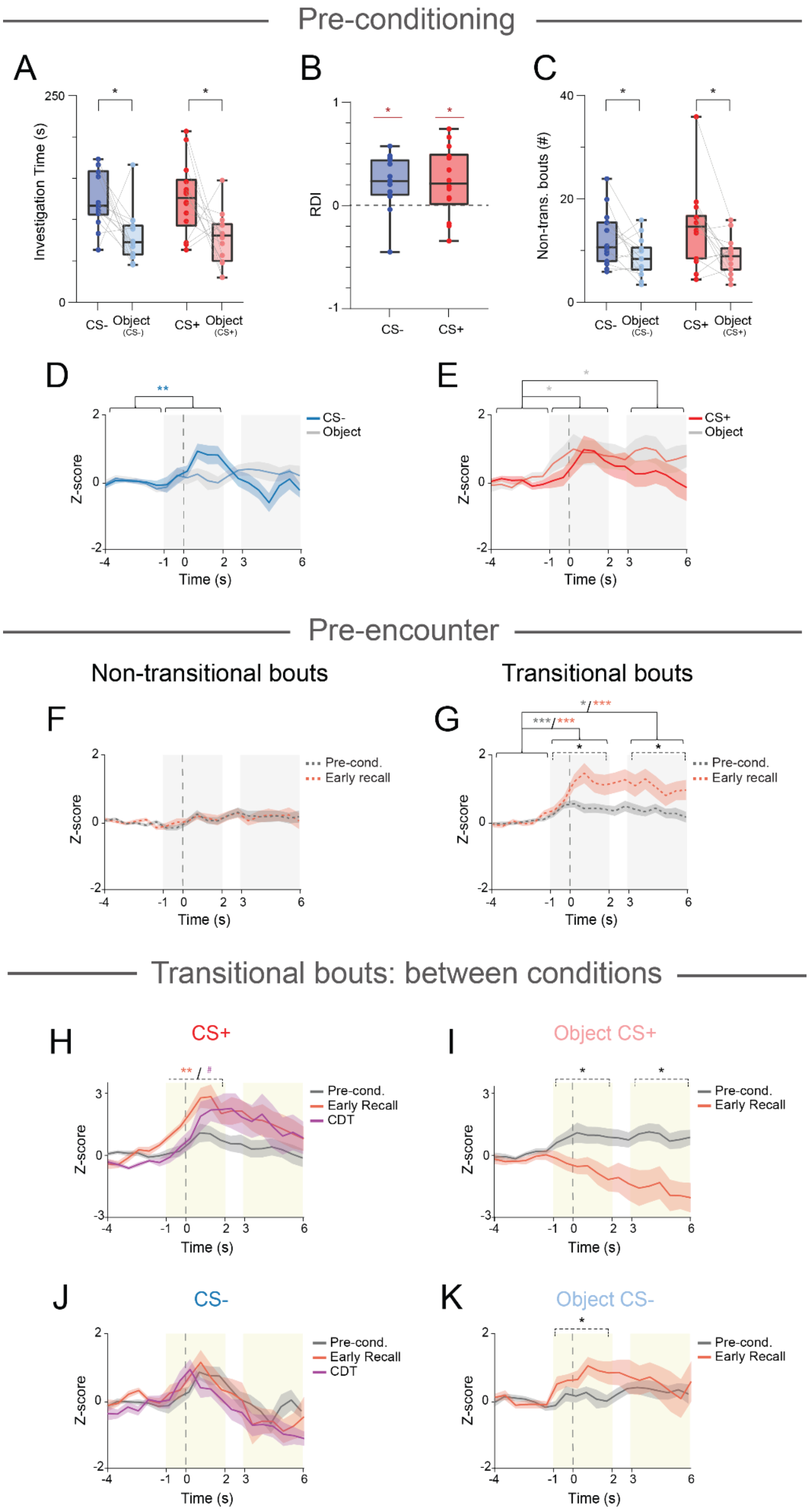

Pre-conditioning
A
B
C
Investigation Time (s)
RDI
Non-trans. bouts (#)
CS-
Object (CS-)
CS+
Object (CS+)
D
E
CS-
CS+
Object
Z-score
Time (s)
Pre-encounter
Non-transitional bouts
Transitional bouts
F
G
Pre-cond.
Early recall
Transitional bouts: between conditions
H
CS+
I
Object CS+
J
CS-
K
Object CS-
Pre-cond.
Early Recall
CDT

**Figure S9. mPFC Pyramidal Neurons show no responses during non-transitional bouts in the social fear conditioning paradigm**

**A.** Median (in box plot) value of mean investigation time of each of the social (C57BL/6J- CS-) and BALB/cJ- CS+) and object stimuli in the 5-min pre-conditioning sessions (n=14 from 14 animals). A significant effect was found for stimulus (*$p<0.01$*) by two-way MM ANOVA.

**B.** Median relative discrimination index (RDI) for the social stimuli in the pre-conditioning with CS+ and CS- sessions.

**C.** Median value for the number of non-transitional bouts to the social (CS+ and CS-) and object stimuli. A significant effect was found for stimulus (*$p<0.01$*) by two-way MM ANOVA.

**D.** Superimposed traces of the mean (±SEM) z-scored calcium signals for transitional bouts averaged across all pre-conditioning sessions with CS- stimuli. Gray-shaded areas represent two distinct time windows of the event, which are used for the statistical analysis. A significant effect was found for time x stimulus interaction (*$p<0.01$)* by two-way MM ANOVA.

**E.** As in **D,** for the pre-conditioning sessions with the CS+ stimuli. A significant effect was found for time (*$p<0.05$)* by two-way MM ANOVA.

**F.** Superimposed traces of the mean (±SEM) z-scored calcium signals shown for non-transitional bouts to empty chambers in the pre-encounter stage of the sessions, averaged across all pre conditioning (in gray) and early recall sessions (in orange). Gray-shaded areas represent two distinct time windows of the event, which are used for the statistical analysis.

**G.** As in **F**, for transitional bouts to empty chambers. A significant effect was found for time *($p<0.001$),* Phase *($p<0.05$)* and time x phase interaction *($p<0.05$)* by two-way MM ANOVA.

**H.** As in **D,** for transitional bouts to the CS+ stimulus. A significant effect was found for time *($p<0.001$)*, stage ($<0.05$), and time x stage interaction (*$p<0.05$)* by two-way MM ANOVA. Yellow-shaded areas represent two distinct time windows of the event, which are used for the statistical analysis. Asterisks on dashed lines represent significance for comparisons between early recall (colored in orange) or conditioned discrimination test (colored in purple) relative to pre-conditioning.

**I.** As in **H**, for transitional bouts to the object presented with the CS+ stimulus. A significant effect was found for stage ($p<0.01$), and time x stage interaction *($p<0.01$)* by two-way MM ANOVA.

**J.** As in **E**, for transitional bouts to the CS- stimulus averaged across the pre-conditioning, early recall, and conditioned discrimination sessions. A significant effect was found for time *($p<0.001$)* by two-way MM ANOVA

**K.** As in **H,** for transitional bouts to the object presented with the CS- stimulus, averaged across the baseline and early recall sessions.

#*$p<0.07$*, *$p<0.05$, **$p<0.01$, ****$p<0.001$*, *post-hoc* paired (within stimulus differences in time) or independent (differences between stimuli or conditions within time windows) t-test with Holm-Šídák correction for multiple comparisons following detection of main effects by ANOVA.

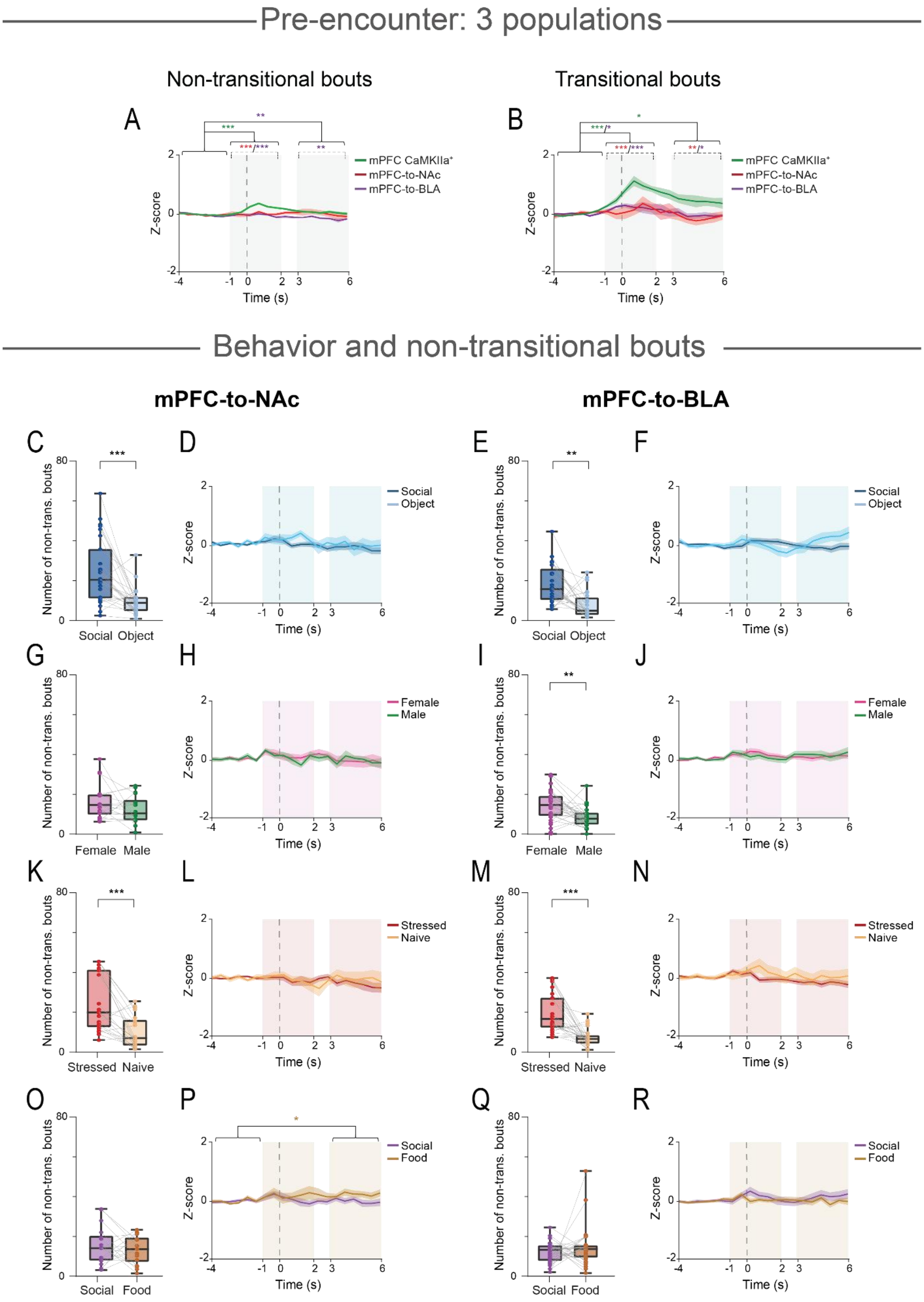
Pre-encounter: 3 populations
Non-transitional bouts
Transitional bouts
A
B
mPFC CaMKIIa+
mPFC-to-NAc
mPFC-to-BLA
Z-score
Time (s)
Behavior and non-transitional bouts
mPFC-to-NAc
mPFC-to-BLA
C
D
E
F
G
H
I
J
K
L
M
N
O
P
Q
R
Number of non-trans. bouts
Social
Object
Female
Male
Stressed
Naive
Food

**Figure S10. mPFC projections to the NAc and BLA are not differentially activated in non-transitional bouts**

**A.** Superimposed traces of the mean (±SEM) z-scored calcium signals shown for non-transitional bouts to empty chambers in the pre-encounter stage of all recorded sessions, averaged across the three recorded neural populations: mPFC CaMKII$\alpha^+$ (in green), mPFC-to-NAc projecting neurons (in red), and mPFC-to-BLA projecting neurons (in purple). Gray-shaded areas represent two distinct time windows of the event, which are used for the statistical Analysis. Asterisks on dashed lines represent significance for comparisons between mPFC-to-NAc projecting neurons (colored in red) or mPFC-to-BLA projecting neurons (colored in purple) relative to mPFC CaMKII$\alpha^+$ population (in green). A significant effect was found for time ($p<0.01$), neural population ($p<0.001$), and time x neural population interaction ($p<0.001$) by two-way MM ANOVA.

**B.** As in **A**, for transitional bouts to empty chambers. A significant effect was found for time ($p<0.001$), neural population ($p<0.001$), and time x neural population interaction ($p<0.001$) by two-way MM ANOVA.

**C.** Median sum of non-transitional bouts to the social stimulus and the object during the 5-minute encounter period of all SP sessions recorded from animals with GCaMP expression in mPFC-to-NAc neuronal cells.

**D.** As in **A**, for non-transitional bouts during the encounter stage with the social stimulus (dark blue) and object (light blue), averaged across all SP sessions recorded from mPFC-to-NAc neuronal cells, and aligned to the bout onset (time '0'). Blue-shaded areas represent two distinct time windows of the event, which were used for the statistical analysis.

**E-F.** As in **C-D**, for SP sessions recorded from mPFC-to-BLA neuronal cells

**G-J.** As in **C-F**, for the SxP task.

**K-N.** As in **C-F**, for the ESPs task.

**O-R.** As in **C-F**, for the SvFP task.

*$**p < 0.001$, $**p < 0.01$,* Wilcoxon signed-rank test in **C**, **E**, **I**, **K**, and **M**.
