## Supplementary material for "Inhibition governs preference encoding in medial prefrontal cortex pyramidal neurons during a binary social choice in mice": Description of Additional Supplementary Files

File Name: Supplementary Movie 1

Description**: Example clip (37s long) from an SP session with optic stimulation delivered during bouts with the social stimulus.**

File Name: Supplementary Movie 2

Description**: Example (31s long) clip from an ESPs session with optic stimulation delivered during bouts with the stressed stimulus.**

File Name: Supplementary Movie 3

Description: **Example (20s long) clip from a SxP session with optic stimulation delivered during bouts with the Female stimulus.** Subject animal still shows repetitive re-engagement with the female stimulus driven by optic stimulation even at lower optic stimulation parameters (4.99mW light power, 7ms pulses at 10Hz).

All three clips demonstrate how optogenetic stimulation of mPFC pyramidal neurons induces an abrupt termination of social interaction, followed by repetitive re-engagement with the same stimulus once the stimulation ceases.
